# Near-infrared light increases functional connectivity with a non-thermal mechanism

**DOI:** 10.1101/459883

**Authors:** Grzegorz M. Dmochowski, Ahmed (Duke) Shereen, Destiny Berisha, Jacek P. Dmochowski

## Abstract

Although techniques for non-invasive brain stimulation are under intense investigation, an approach that has received limited attention is *transcranial photobiomodulation* (tPBM), the delivery of near-infrared light to the brain with a laser directed at the scalp. Here we employed functional magnetic resonance imaging (fMRI) to measure the Blood-Oxygenation-Level Dependent (BOLD) signal in *n* = 20 healthy humans while concurrently stimulating their right frontal pole with a near-infrared laser. We failed to detect an evoked BOLD response at illumination. However, functional connectivity with the illuminated region increased by an average of 10% during stimulation, with some connections strengthening by as much as 40%. 23% of connections with the illuminated region experienced a significant acute increase, with the time course of connectivity exhibiting a sharp rise at illumination onset. Brain-wide connectivity increases were also observed, with connections in the stimulated hemisphere showing a significantly larger increase than those in the non-stimulated hemisphere. We subsequently employed MR Thermometry to measure brain temperature during tPBM (separate cohort, *n* = 20), and found no significant temperature differences between active and sham stimulation. Our findings suggest that near-infrared light synchronizes brain activity with a non-thermal mechanism, underscoring the promise of tPBM as a new technique for stimulating brain function.

## Introduction

Causal interventions of neural activity are necessary to identify the brain circuits underlying complex behaviors, and also to spur the development of non-systemic therapies for neurological and psychiatric disorders. The most popular approaches employ mild electric currents, magnetic fields (Dayan et al., 2013), or ultrasonic waves (Naor et al., 2016). These techniques are aimed at directly evoking neuronal activation or modulating cortical excitability. In contrast, a more recent neuromodulation technique employs light to target the brain’s energy metabolism pathway. Termed *transcranial photobiomodulation* (tPBM), this relatively unknown approach delivers near-infrared light to the brain via transcranial transmission with a laser or light emitting diode (LED) (Hamblin, 2016). The purported mechanism of action in tPBM is the absorption of light by cytochrome *c* oxidase (CCO), the terminal enzyme in the mitochondrial electron transport chain, leading to increased energy metabolism (Cassano et al., 2016). Findings from small animal models suggest that tPBM increases cerebral blood flow (CBF) (Uozumi et al., 2010) and cortical ATP (Mochizuki-Oda et al., 2002), while reducing reducing inflammatory markers (Moreira et al., 2009; Zhang et al., 2014) and apoptosis (Wong-Riley et al., 2005; Yu et al., 2015; Salehpour et al., 2017). tPBM has also been reported to ameliorate amyloid beta accompanying neurodegeneration (Lu et al., 2017) and oxidative damage from sleep deprivation (Salehpour et al., 2018). However, it is largely unknown whether such effects could be achieved in human.

It is important to determine whether tPBM represents a viable form of non-invasive brain stimulation, and if so, through what mechanism of action. To date, there are limited reports of the neurophysiological effects of tPBM in the human brain. Tian et al. (2016) employed near-infrared resonance spectroscopy (NIRS) and reported increased cerebral oxygenation in both hemispheres during tPBM. A recent report suggests that tPBM increases the power of electrophysiological oscillations as measured by the scalp electroencephalogram (EEG) (Wang et al., 2019). Behavioral investigations have reported that tPBM may improve performance on cognitive tests of working memory and attention (Barrett and Gonzalez-Lima, 2013; Hwang et al., 2016; Blanco et al., 2017b, a). The ability of tPBM to accelerate cerebral energy metabolism in humans remains largely untested: Does the human brain respond metabolically to tPBM, and if so, with what temporal dynamic? Moreover, given that tPBM involves depositing energy into the brain, it is critical to ascertain whether heating is involved in any observed effects.

Functional magnetic resonance imaging (fMRI) affords the opportunity to test the merit of tPBM in the human brain. In particular, the blood-oxygenation-level dependent (BOLD) signal (Ogawa et al., 1990) is closely linked to energy metabolism, with both cerebral blood flow (CBF) and the cerebral metabolic rate of oxygen (CMRO2) contributing to the measured signal (Buxton, 2013). Traditionally, BOLD has been employed as an indirect measure of neural activation via neurovascular coupling. In addition, MR Thermometry (Rieke and Butts Pauly, 2008) permits the measurement of brain tissue temperature during stimulation, such that the presence and magnitude of heating can be determined. Despite this, the physiological effects of tPBM have only scarcely been probed with MRI, with recent reports indicating reductions in somatosensory evoked activity of healthy subjects (El Khoury et al., 2019), and altered functional connectivity in a cohort of six chronic stroke patients (Naeser et al., 2019).

Here we measured the neurophysiological response of the human brain to tPBM with two separate MRI techniques. We first recruited *n* = 20 healthy participants to receive a 10-minute application of tPBM to the right frontal pole while having their hemodynamic activity recorded with BOLD-fMRI. Contrary to our expectations, we did not detect an evoked BOLD response at illumination. However, we found a robust effect on functional connectivity, with the illuminated region showing a mean 10% increase relative to the pre-stimulation period, with some connections strengthened by as much as 40%. The time course of connectivity exhibited a sharp increase at illumination onset. We also found enhanced connectivity between regions outside of the illuminated area, with a significantly larger increase in the stimulated hemisphere. We subsequently employed MR thermometry to measure brain temperature changes during tPBM with the same parameters as the BOLD study (separate cohort of *n* = 20). We failed to detect a thermal effect of tPBM, with the temperature in the illuminated region not varying significantly from sham stimulation at any time points.

## Materials and Methods

### Subjects

All experimental procedures were approved by the Institutional Review Board of the City University of New York. We recruited *N* = 40 participants (20 females) from the local New York City population. In an attempt to achieve uniform baseline measures of CBF and CMRO2 in our sample, only subjects aged 18-40 were considered: the cerebral metabolic rate of oxygen (CMRO_2_) has been found to decrease monotonically with age (Yamaguchi et al., 1986; Marchal et al., 1992). The mean age of the participants was 24.8 ± 4.6 years. During recruitment, we employed those exclusionary criteria common to MRI (e.g., patients with cardiac pacemakers, neurostimulation systems, or claustrophobia were excluded). All subjects completed the experiments and there were no major adverse effects. In the BOLD-fMRI study, one subject complained of a headache following the scan, which may have been caused by the headgear that was worn throughout the (~1 h) experiment.

### Experimental Design

*BOLD-fMRI Study. N* = 20 subjects (10 females) participated. The BOLD signal was continuously acquired for 30 minutes. The laser was turned on 10 minutes after the onset of the scan and remained active for a 10-minute duration. Subjects were not made aware of when the laser was turned on or for how long. From verbal post-experimental surveys, subjects did not perceive any sensations (thermal or otherwise) during illumination. *MR Thermometry Study. N* = 20 subjects (10 females) participated. All subjects performed two sessions in succession. Throughout each 20.5-minute session, brain temperature was measured with a temperature-sensitive MRI sequence. In one session (“active”), the laser was turned on 172 s (21 TRs) into the scan and remained on for 10 minutes. In the other session (“sham”), the laser remained off throughout. As the switch controlling the laser was housed in the MRI control room, subjects could not see or hear the operation of the device. The order of sham and active sessions was randomized and counterbalanced across subjects.

### Transcranial Photobiomodulation

tPBM was applied at a wavelength of 808 nm, selected based on a previous study that demonstrated an improvement in cognitive task performance after tPBM in healthy volunteers (Barrett and Gonzalez-Lima, 2013). This wavelength falls within the so-called “optical window” in which absorption by common chromophores (water, melanin, and hemoglobin) is low (Hamblin and Demidova, 2006), allowing for deeper penetration through the human scalp and skull and into the brain as compared to other candidate wavelengths (Tedford et al., 2015; Pitzschke et al., 2015). A class IV 10W diode laser (Ultralasers MDL-N-808-10000) powered by a laser driver (Ultralasers PSU-H-LED) provided the monochromatic light. A calibrated photodetector was employed to set the laser power to 250 mW prior to each experiment. Given a 1 cm diameter aperture in the headgear worn by participants, this resulted in an intensity of 318 mW/cm^2^, which is within the ANSI safety limit for human tissue (330 mW/cm^2^). As the duration of stimulation was set to 10 minutes, the total incident light energy was 150 J. It should be noted that the coherent nature of the laser is not believed to contribute to the action of tPBM. A given photon undergoes multiple scattering events on its path to the brain, thereby losing its original phase information. Light emitting diodes (LEDs) have elicited effects in previous PBM experiments (Chung et al., 2012). A laser was employed in this case to facilitate light delivery at the desired power.

The laser output was coupled directly into a custom made, multimode optical fiber (Thorlabs FT400EMT) whose core diameter was 400 μm. In order to ensure MR compatibility, the distal end of the optical fiber was fitted with a ceramic ferrule that was affixed to a custom 3D-printed headgear worn by participants. The headgear, measuring 5.5 cm × 3.2 cm × 2.7 cm, contained a clamp which secured the ferrule. The headgear was secured against the subject’s head such that its aperture was flush against the forehead. The aperture was centered at location “Fp2” (right frontal pole) of the 10/20 standard system for electroencephalography (Jasper, 1958), where each participant’s Fp2 location was measured and marked on the scalp prior to the experiment. This location matched that employed previously (Barrett and Gonzalez-Lima, 2013). Note that stimulating the forehead yields better penetration through the scalp due to the absence of scattering by the hair. For all but two subjects, Vitamin E markers were placed on the headgear so that the location of light incidence could be registered with the anatomical MRI. All subjects wore protective goggles in addition to the laser headgear throughout all experiments.

### Estimating the Illuminated Region

The primary region-of-interest (ROI), referred to herein as the “illuminated region”, was constructed based on numerical simulations of light propagation through the human head. We employed the Monte Carlo Multi-Layered (MCML) software (Wang et al., 1995), which models an infinitely narrow photon beam normally-incident on multiple layers of turbid material. The four layers here corresponded to scalp, skull, cerebrospinal fluid (CSF), and brain. The properties of the model are provided in Table S1. The output of the simulation was the light absorption (J/cm^3^) in the volume, computed over a discrete grid in cylindrical coordinate space. To define the illuminated region, we computed the smallest cylinder which contained at least 99% of the total absorption. An exhaustive search procedure produced a cylindrical ROI with a radial extent of 2.1 cm and axial extent of 3.9 cm. This ROI was “projected” onto each subject’s anatomical MRI from the position of the aperture.

### MRI, fMRI, and MR Thermometry

*BOLD-fMRI.* Imaging was performed with a Siemens Magnetom Skyra 3 Tesla scanner. A 16-channel transmit/receive head coil was used for data acquisition. Structural images were acquired with a T1-weighted MPRAGE sequence (FOV 230 mm, in plane resolution 256 x 256, 224 slices with a thickness of 0.9 mm, TI = 1000 ms). Functional BOLD scans were acquired with a multi-echo EPI sequence (FOV 228 mm, in plane resolution 90-by-90, 60 slices with a thickness of 2.5 mm, TR=2800 ms, Flip Angle = 82 degrees). The three echo times were 12.8 ms, 34.3 ms, and 55.6 ms, which allowed for the characterization of the T2* decay of the BOLD signal (Posse et al., 1999). The duration of the BOLD scans was 30 minutes (645 volumes). Subjects were instructed to rest but stay awake and to not think about anything in particular. *MR Thermometry*. Imaging was performed with a Siemens Prisma 3 Tesla scanner. Signal excitation was done with the built-in body coil, and a 20-channel phased array head/neck coil was used for data acquisition. Structural images were acquired with a T1-weighted MPRAGE sequence (FOV 256 mm in read and 240 mm in phase-encode directions, in-plane matrix size of 256 x 240, 208 sagittal slices with a thickness of 1mm, TR/TE/TI = 2400/2.15/1000ms, flip angle = 8 deg, 2x GRAPPA acceleration factor, fat suppression using fast water excitation, and a total acquisition time of 5:26 min.). To measure brain temperature changes during tPBM, we employed a custom 3D EPI phase-difference imaging sequence that exploits the temperature dependence of the Proton Resonance Frequency (PRF) (Ishihara et al., 1995; Rieke and Butts Pauly, 2008). Baseline phase was first measured as an average over 3 initial frames, and subsequent image phases were referenced to this baseline phase. The phase difference has a linear dependence on the temperature change from baseline (Rieke and Butts Pauly, 2008). Thermometry scans covered a FOV of 192 mm in both read and phase directions, an in-plane matrix size of 64-by-64, with 32 interleaved slices per slab of thickness of 3 mm, TR/TE=25/17 ms, Flip Angle = 10 degrees, EPI factor = 7, echo spacing = 0.93 ms, bandwidth = 1302 Hz/Px, with no fat suppression or accelerated imaging used. The duration of the scans was 20.5 minutes (150 volumes).

### BOLD Preprocessing

Preprocessing of BOLD data was performed with the AfNI software package (Version 17.3.03) (Cox, 1996), scripted in the Matlab programming language (Mathworks, Natick, MA).

The anatomical image was first skull-stripped using the *3dSkullStrip* function, whose output was then used to create a brain mask via the *3dAutomask* function. The anatomical image was segmented to produce tissue masks for the CSF, grey matter, and white matter via the *3dSeg* routine. The first three frames of all BOLD series were excluded from analysis. The function *3dDespike* was applied to the raw BOLD series to remove large transients. Slices were aligned to the onset of each TR. Motion correction was performed by aligning each volume of the BOLD series to a reference volume (i.e., frame 3). We aligned the BOLD data to the corresponding anatomical image, and applied a non-linear warping procedure to then transform the data to the Talairach coordinate space. Spatial smoothing with a full-width-half-max (FWHM) of 4 mm was then performed. Each voxel’s time series was normalized to a mean of 100. The time series of the motion alignment parameters and their derivatives were linearly regressed out of the data. The BOLD was then band-pass filtered to the range 0.01 – 0.1 Hz. Volumes during which the derivative of the motion parameters exceeded a norm of 0.3 were censored. Volumes during which more than 15% of the voxels were identified as outliers, defined as those samples exceeding 5.8 times the median absolute deviation, were also censored.

### Event-related BOLD analysis

To probe evoked (i.e., time-locked) changes in the BOLD signal at illumination, we aligned the BOLD series of all subjects such that the precise onset TR was consistent across the group. We computed the mean and standard error of the group-averaged BOLD for the illuminated region and two other ROIs in the vicinity of illumination: the right orbital gyrus, and the right middle frontal gyrus. BOLD time series were first averaged within each ROI and then across subjects. To test for a significant evoked response, we formed surrogate data records following the approach of Theiler et al. (1992), where the BOLD series at every voxel and echo had its phase spectrum permuted while preserving its amplitude spectrum. Importantly, within each subject, the same permutation was applied to all voxels, such that the spatial correlation of the data were preserved. Note that the temporal autocorrelation of the BOLD signals is also maintained after phase randomization. The procedure was repeated 500 times such that a null distribution of group-averaged BOLD could be constructed at every time point during and after illumination. The true group-averaged BOLD was then compared to the sample null distribution values, with the p-value following as the proportion of permutations exceeding the true BOLD amplitude at a given time point. We corrected the resulting p-values for multiple comparisons by controlling the FDR at 0.05.

### Functional Connectivity Analysis

Functional connectivity was performed on time series from 151 ROIs formed by the union of the Destrieux atlas (75 ROIs in each hemisphere) (Destrieux et al., 2010) and the illuminated region. The time series of grey matter voxels comprising each ROI were averaged prior to connectivity analysis. As we observed a transient increase in functional connectivity during the first five minutes of recording, we retained only the second half of the pre-illumination period for analysis (Figure S1). In other words, pre-illumination values were measured once the data reached a stable level. For each of the three experimental time segments: before illumination (5 minutes), during illumination (10 minutes), and immediately following illumination (10 minutes), we measured functional connectivity as the Pearson correlation coefficient between each unique pair of ROI time courses. Correlation coefficients were Fisher transformed prior to all statistical tests. For each participant and echo, we also computed the *total connectivity* of the illuminated region as the mean correlation between the illuminated region and each of its 150 connecting ROIs (mean across 150 connecting ROIs).

We tested for significant effects of tPBM on functional connectivity with the permutation testing approach advocated in the field of dynamic functional connectivity (dFC) (Preti et al., 2017; Handwerker et al., 2012). Surrogate data records were constructed by permuting the phase spectrum of voxel time series, in all cases preserving spatial correlations among voxels and autocorrelations within time series. From this, we were able to construct null distributions of the group-level difference in functional connectivity between the illumination and pre-illumination period, and also between the post-illumination and pre-illumination period. We also employed the permutation test approach when testing for changes in total connectivity and brain-wide connectivity. In other words, we performed the same sequence of operations as was carried out on the actual data but on the permuted data. In all cases, we corrected for multiple comparisons by controlling the FDR at 0.05. When testing for the presence of significantly larger effects in the right hemisphere, we performed paired, two-sample t-tests (*n* = 20) on Fisher transformed correlation coefficients that were averaged across all connections within the corresponding region.

### MR Thermometry Preprocessing

Brain temperature recordings followed a similar preprocessing procedure as the BOLD signals. The AfNI function *align_epi_anat.py* was employed to perform the combined operations of slice time correction, motion correction, and registration of temperature volumes to the anatomical images. We used the *lpa* cost function and registered the magnitude of the PRF to define the transformation which was then applied to the phase data. Phase difference images were smoothed with a FWHM of 8 mm using *3dBlurInMask.* We employed the PRF equation to convert the phase differences to temperature changes:

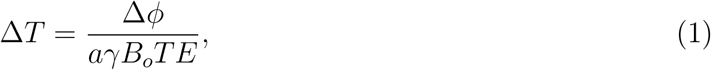

where Δ*T* (units of °C) is the temperature change from baseline, Δ*ϕ* (units of radians) is the phase difference from baseline, *a* = −0.01 × 10^-6^ is the PRF constant modeling the sensitivity of the resonance frequency to temperature changes, *B*_0_ = 3 T, *TE* = 0.017 s, *γ* = 2π × 42.58 × 10^6^ rad/s. We then regressed out the time series of the *B_o_* reference as well as the time series of the motion parameters and their derivatives. Voxels whose mean signal power was greater than 4 standard deviations above the mean of all voxels were censored.

### MR Thermometry Analysis

In order to ensure the accuracy of the MR Thermometry sequence and subsequent data analysis, we conducted recordings with an agar phantom that was heated up with the laser at a high intensity. Ground-truth temperature in the phantom was simultaneously measured with an infrared temperature sensor and used to validate the MR-derived temperature.

To test for the presence of heating during the human tPBM experiments, we averaged the temperature change (from baseline) across all voxels in the illuminated region, here including the white matter. This was computed separately for all time points and both active and sham conditions. We then performed paired two-tailed Wilcoxon sign rank tests with *n* = 20 to detect time points during which there was a significant difference in temperature change between active and sham stimulation. Correction for multiple comparisons was implemented by controlling the FDR at 0.05.

## Results

To investigate the effect of tPBM on hemodynamic activity and temperature in the human brain, we conducted experiments combining laser stimulation with MRI in healthy human participants. tPBM was applied to the right frontal pole (scalp location Fp2) of *n* = 20 healthy participants with a monochromatic 808 nm laser at an intensity of 318 mW/cm^2^ and a 10-minute duration. A multi-echo (13 ms, 34 ms, 55 ms) BOLD-fMRI sequence was employed to capture hemodynamic changes due to tPBM, with a 25-minute analysis window considering the 5 minutes leading up to illumination, the 10-minute stimulation period, and the 10 minutes immediately after illumination (Figure 1). To determine whether tPBM produced significant heating of the brain, we then employed MR Thermometry to measure brain temperature in a separate cohort of *n* = 20 participants receiving tPBM with the same dose as the BOLD-fMRI study.

**Figure 1:**
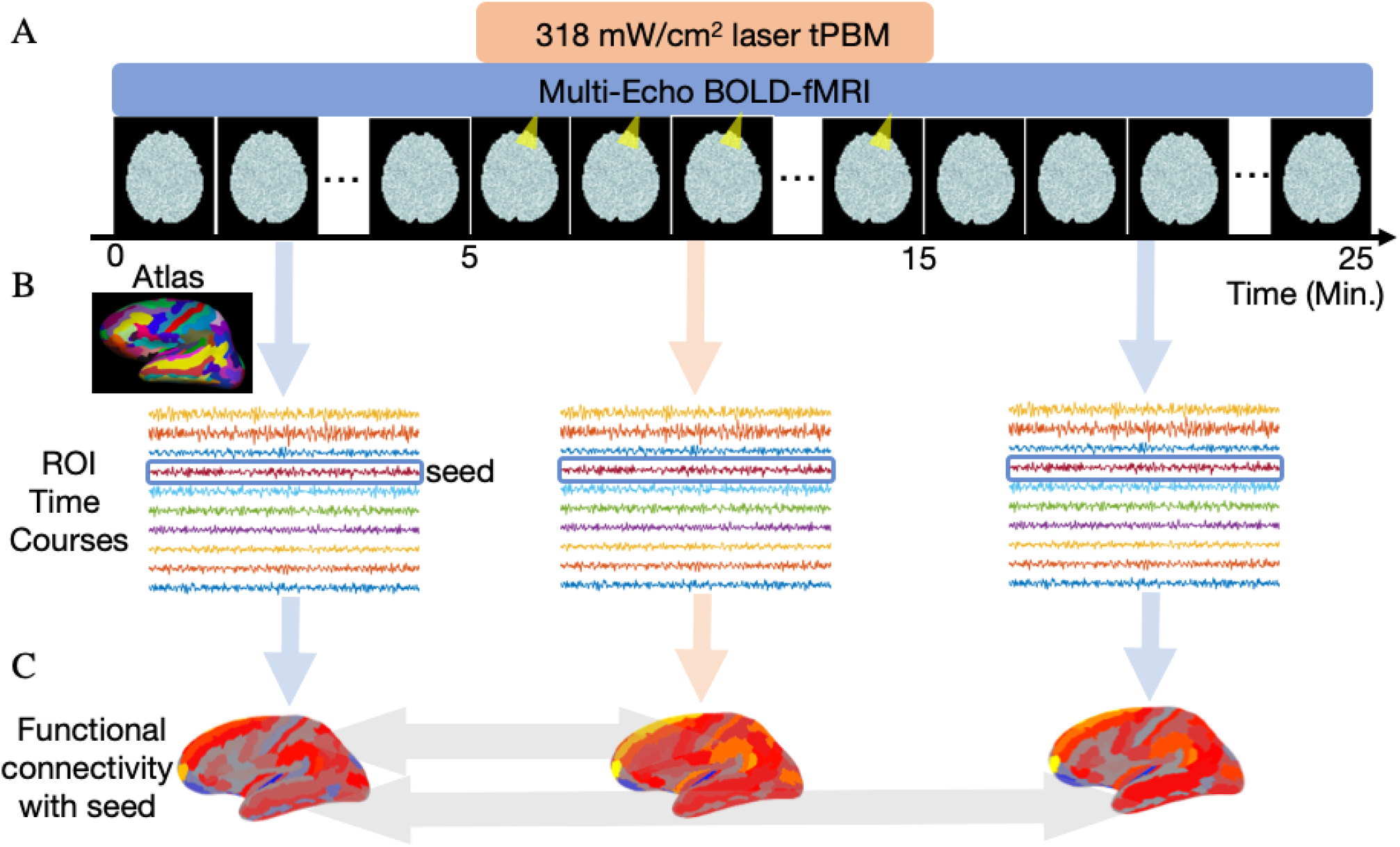
Experimental Design. (**A**) To identify the effect of tPBM on hemodynamic activity in the human brain, we recorded the BOLD-fMRI signal from *n* = 20 healthy participants before, during, and after illumination. tPBM was applied with an 808 nm laser at an intensity of 318 mW/cm^2^ beginning 5 minutes after the start of the 25-minute analysis window. The duration of tPBM was 10 minutes. (**B**) BOLD signals were converted into region-of-interest (ROI) time series corresponding to 151 cortical regions from the Destrieux atlas (Destrieux et al., 2010) and the illuminated region. (**C**) Functional connectivity was computed as the Pearson correlation between the time courses of a “seed” region and connecting ROIs. We measured the difference in functional connectivity between the illumination and pre-illumination periods, and also between the post-illumination and pre-illumination periods (indicated with light gray arrows). Permutation tests were conducted to test for significant changes in functional connectivity, with multiple comparisons corrected by controlling the false discovery rate (FDR) at 0.05.

### No effect on evoked BOLD response

An individualized region-of-interest (ROI) was defined for each subject based on the precise scalp site of illumination and a simple model of transcranial light propagation (see *Methods*). The cylindrical ROI extended approximately 2 cm radially and 4 cm axially, designed to contain at least 99% of the total light absorption. In what follows, we refer to this ROI as the “illuminated region”.

We first probed changes to the amplitude of the BOLD signal evoked by tPBM. Due to the purported action of near-infrared light on mitochondrial activity, we expected to see a time-locked hemodynamic change near the site of illumination. Against our expectation, we did not find a significant BOLD change during or immediately after illumination at any of the 3 echos (*p* > 0.05 for all time points, permutation test employing phase-randomized surrogate data, Figure 2A). We also probed changes in nearby atlas-derived ROIs such as the right orbital frontal gyrus and right middle frontal gyrus. We failed to detect a significant BOLD change at any time points for these neighboring ROIs (*p* > 0.05 for all time points, permutation test, Figure 2B-C).

**Figure 2:**
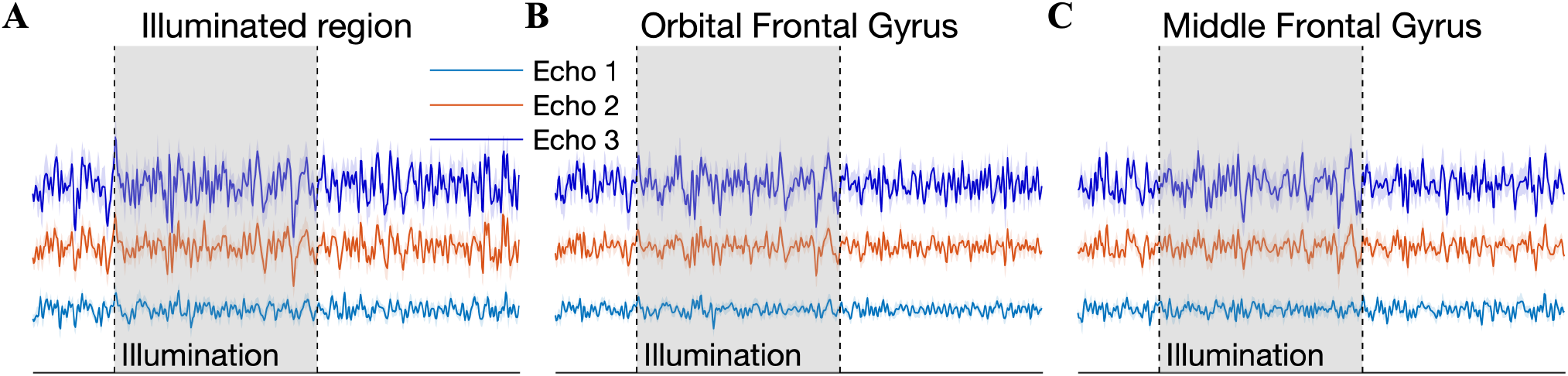
No effect of tPBM on evoked BOLD response. (**A**) Group-averaged time courses of the BOLD signal in the illuminated region (right frontal pole) before, during, and after illumination. All three echos of the T2* signal are shown, where the echo times are 13 ms, 34 ms, and 55 ms. BOLD signals of the multiple subjects were aligned to illumination onset prior to averaging. There were no time points during or after illumination at which the BOLD exhibited significant deviation from the null distribution (*p* > 0.05 for all time points, permutation test employing phase randomized surrogate data). (**B**) Same as (A) but now for the right orbital frontal gyrus. No significant differences were detected. (**C**) There was also no significant deviation from the null distribution in the right middle frontal gyrus. Thus, we found no evidence that tPBM evoked a BOLD response in the vicinity of the illuminated region.

### Near-infrared light increases functional connectivity with the illuminated region

In the resting state, BOLD has most often been investigated in the context of functional connectivity: the temporal correlation of the hemodynamic signal across different brain regions (Fox and Raichle, 2007). To account for the possibility of tPBM affecting phase (as opposed to amplitude), and to probe possible “network” effects on hemodynamic activity, we measured functional connectivity before, during, and after illumination. Connectivity analysis was carried out on a set of 151 ROIs from the Destrieux brain atlas (75 cortical ROIs in each hemisphere) (Destrieux et al., 2010), and the illuminated region.

We first compared functional connectivity between the illuminated region and all other ROIs across the three temporal segments of the experiment. During illumination, a robust increase in connectivity was observed at all echo times (results for Echo 3 shown Figure 3A-B). Functional connectivity was visibly increased in the frontal, temporal, and parietal cortex of both hemispheres. Similar increases in functional connectivity were also found at Echos 2 and 3 (Figures S2–S3). Of the 150 connections that the illuminated region made with other regions, 34(17 significant connections with the left hemisphere, 17 with the right) exhibited a statistically significant increase during illumination relative to the pre-stimulation period (permutation test with phase-randomized surrogate data records modeling static connectivity, corrected for multiple comparisons with the false discovery rate at 0.05; shown for Echo 3 in Figure 3C-D). At echos 2 and 3, the number of significantly enhanced connections was 11 and 11, respectively. Averaged across all connections, the percent increases in functional connectivity were 9.68 ± 4.22%, 10.60 ± 4.38%, and 12.67 ± 5.09% at echos 1, 2, and 3, respectively (means ± SEMs across *n* = 20 subjects). The largest measured increase was with the left postcentral sulcus (parietal lobe; 40% increase at echo 2). At Echo 3, the significant increases in functional connectivity were with connections to the left and right cingulate, left and right superior parietal cortex, left and right precuneus, among others. The functional connectivity with the illuminated region for all connecting ROIs and echos is listed in Tables S2–S3. Only a few connections passed significance in the post-illumination period: the left subparietal sulcus (echo 1), the left and right cingulate gyri (echo 1), and the left precentral sulcus (echo 3).

**Figure 3:**
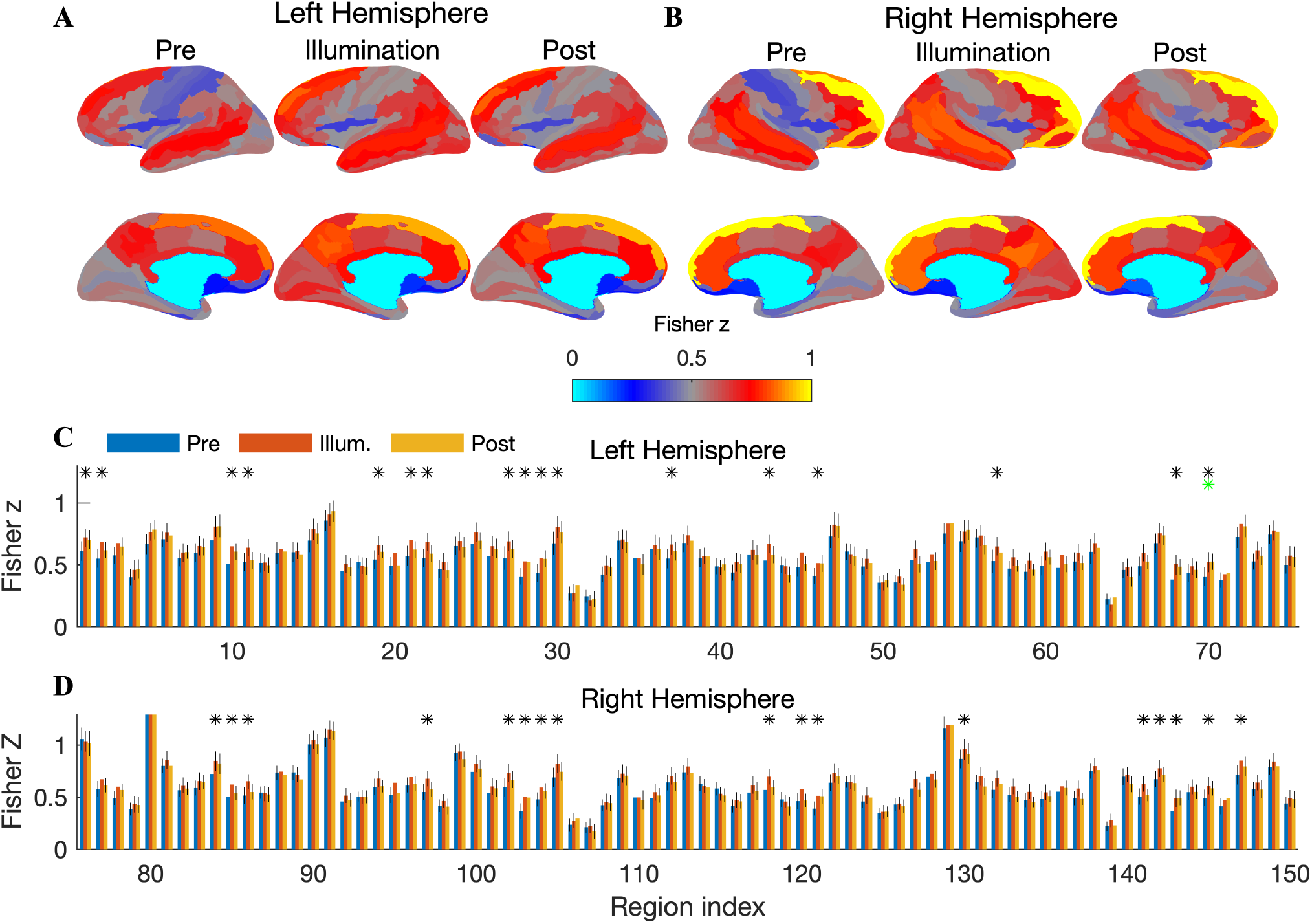
Increased functional connectivity with the illuminated region during tPBM. (**A**) Cortical surfaces display the group-averaged functional connectivity (Fisher transformed Pearson correlations) between the illuminated region and all left hemispheric ROIs, shown separately for the pre-illumination, illumination, and post-illumination periods. During tPBM, increased connectivity was apparent in the dorsal frontal cortex, the temporal lobe, and the parietal cortex. Note that the magnitude of the increase is reduced following illumination. (**B**) Same as A but showing connectivity with the right hemisphere. Increased connectivity with the frontal, temporal, and parietal cortices was readily observed. (**C**) Bar graphs show Fisher transformed correlation coefficients between the illuminated region and each cortical region in the left hemisphere. Error bars depict the SEM across *n* = 20 subjects. Connections that exhibited statistically significant increases during illumination are denoted with a black asterisk (permutation test, corrected for multiple comparisons by controlling the FDR at 0.05). 17 of the 75 connections (23%) showed a significant increase. Green asterisks denote a significant post-illumination effect, which here was only observed with the left precentral gyrus (frontal cortex). (**D**) Same as C but now for the right hemisphere. 17 of the 75 connections exhibited a significant increase during illumination. ROI labels for all cortical regions in the bar graphs are provided in Tables S2–S3, where functional connectivity before, during, and after illumination is listed for all echos.

To further probe the observed increase in functional connectivity with the illuminated region, we measured the time course of connectivity by employing a sliding window of length 1 minute with a 75% overlap among successive windows. To minimize the number of statistical comparisons, here we considered *total connectivity*, which we define as the mean correlation between the illuminated region and its connections (i.e., mean across 150 connecting ROIs). We observed a sharp increase in connectivity at illumination onset, particularly at echo 1 (Figure 4A). Peak connectivity occurred 5 minutes after illumination onset for echo 1, 7 minutes after illumination onset for echo 2, and 2.5 minutes after illumination onset for echo 3. The distinct time courses at the three echos suggest that illumination may have differentially modulated CBF and CMRO2 (see *Discussion*).

**Figure 4:**
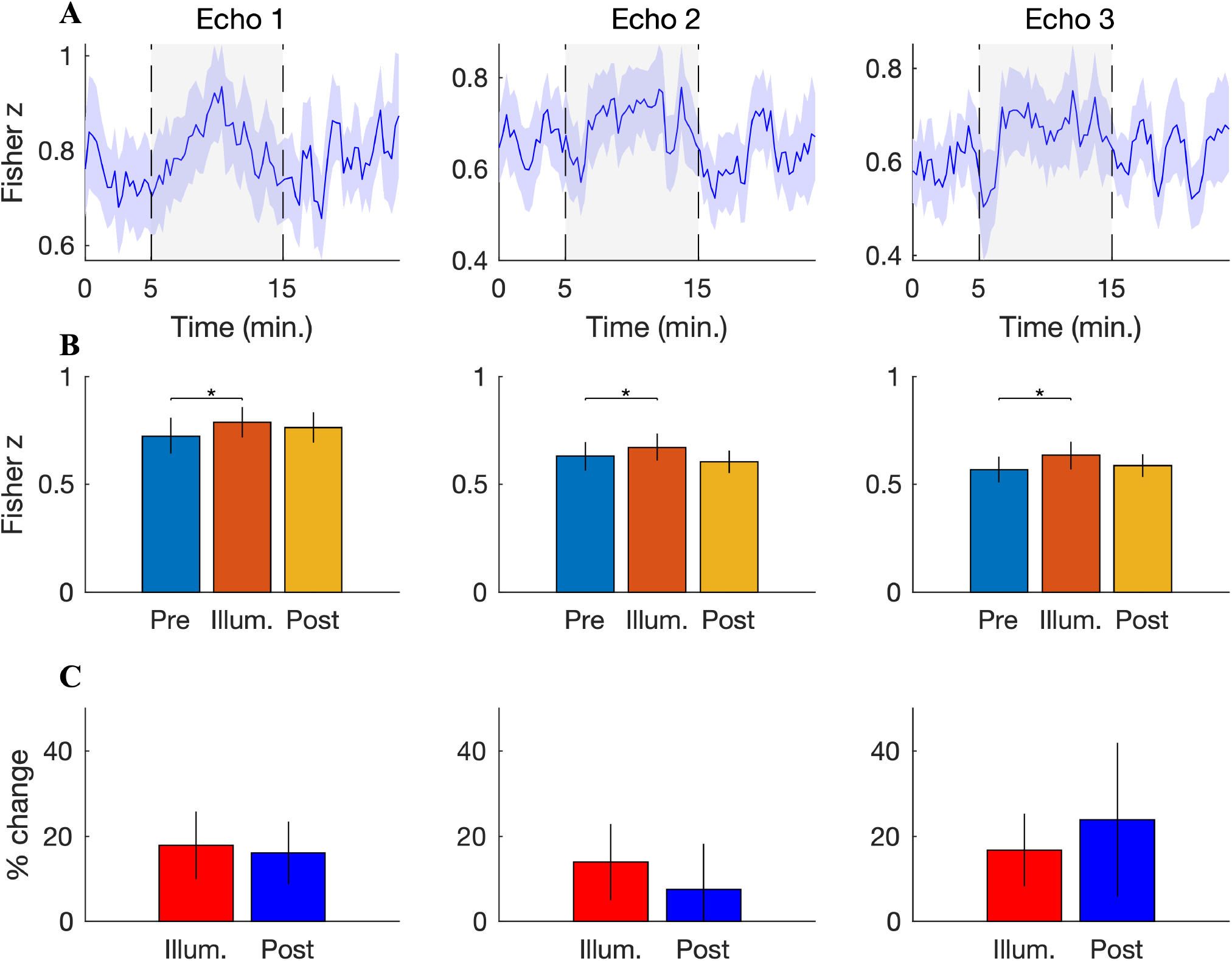
Total connectivity with the illuminated region increases during stimulation. (**A**) The time course of functional connectivity with the illuminated region, computed with a sliding window of one minute duration and 75% overlap. Functional connectivity was averaged first across all 150 connecting ROIs and then across subjects (shaded error bars show SEM across *n* = 20 subjects). At Echo 1 (left), a sharp increase in connectivity was observed at illumination onset, peaking approximately 5 minutes into the stimulation window. The connectivity time courses of Echos 2 (middle) and 3 (right) exhibited a brief negative dip at light onset, followed by a high connectivity period that was sustained throughout illumination. (**B**) Grand mean functional connectivity with the illuminated region before, during, and after illumination. Functional connectivity was averaged across the 150 connecting ROIs, time, and subjects, with the error bars denoting SEM across subjects. A significant increase during illumination was resolved for all echos (permutation test, *p* = 0.018, *p* = 0.050, and *p* = 0.008 for Echos 1, 2, and 3, respectively). (**C**) During illumination, mean increases in total connectivity of 18%, 14%, and 17% were observed during Echos 1 (left), 2 (middle), and 3 (right), respectively. After illumination, the mean percent changes were 16%, 8%, and 24%. Error bars denote the SEM of the percent change in functional connectivity, which was computed first for each subject and then averaged to arrive at the depicted values.

For each of three temporal segments (before, during, and after illumination), we measured total connectivity but now averaging across the entire segment duration. During illumination, we found a significant increase at all echos (permutation test, *p* = 0.018, *p* = 0.050, and *p* = 0.008 for echos 1, 2, and 3, respectively; Figure 4B). We were not able to resolve a significant postillumination effect on total connectivity (p > 0.05 for all echos; Figure 4B). The mean percent change in total connectivity during the illumination period was 17.9 ± 7.9 %, 14.0 ± 8.9 %, and 16.8 ± 8.5 % for echos 1, 2, and 3, respectively (means ± SEMs across *n* = 20 subjects; Figure 4C). After illumination, the mean percent changes were 16.1 ±7.3 %, 7.7± 10.7 %, and 23.9± 18.0 % for echos 1, 2, and 3, respectively (Figure 4C).

### Brain-wide increases in functional connectivity

In order to probe the potential effects of tPBM on connectivity between brain areas outside of the illuminated region, we measured correlation matrices encompassing all connections among the 151 ROIs for the three segments of the experiment (shown for Echo 3 in Figure 5A-C). The illumination period was marked by a pronounced connectivity increase at numerous segments of the correlation matrix, in particular the three quadrants corresponding to connections with ROIs in the stimulated right hemisphere (Figure 5D). Notice the relative absence of increases in the lower-left quadrant, which corresponds to connections within the left hemisphere. In all, 219 connections exhibited statistically significant increases during the illumination period at Echo 3. 111 and 198 were found at Echos 1 and 2, respectively (Figure S4–S5). The increase was largely dampened following illumination (Figure 5E). Nevertheless, we did detect a small number of connections (i.e., 27 at Echo 3) exhibiting a significant increase after illumination (Figure 5E). There were 37 and 9 significant post-illumination connections detected at echos 1 and 2, respectively.

**Figure 5:**
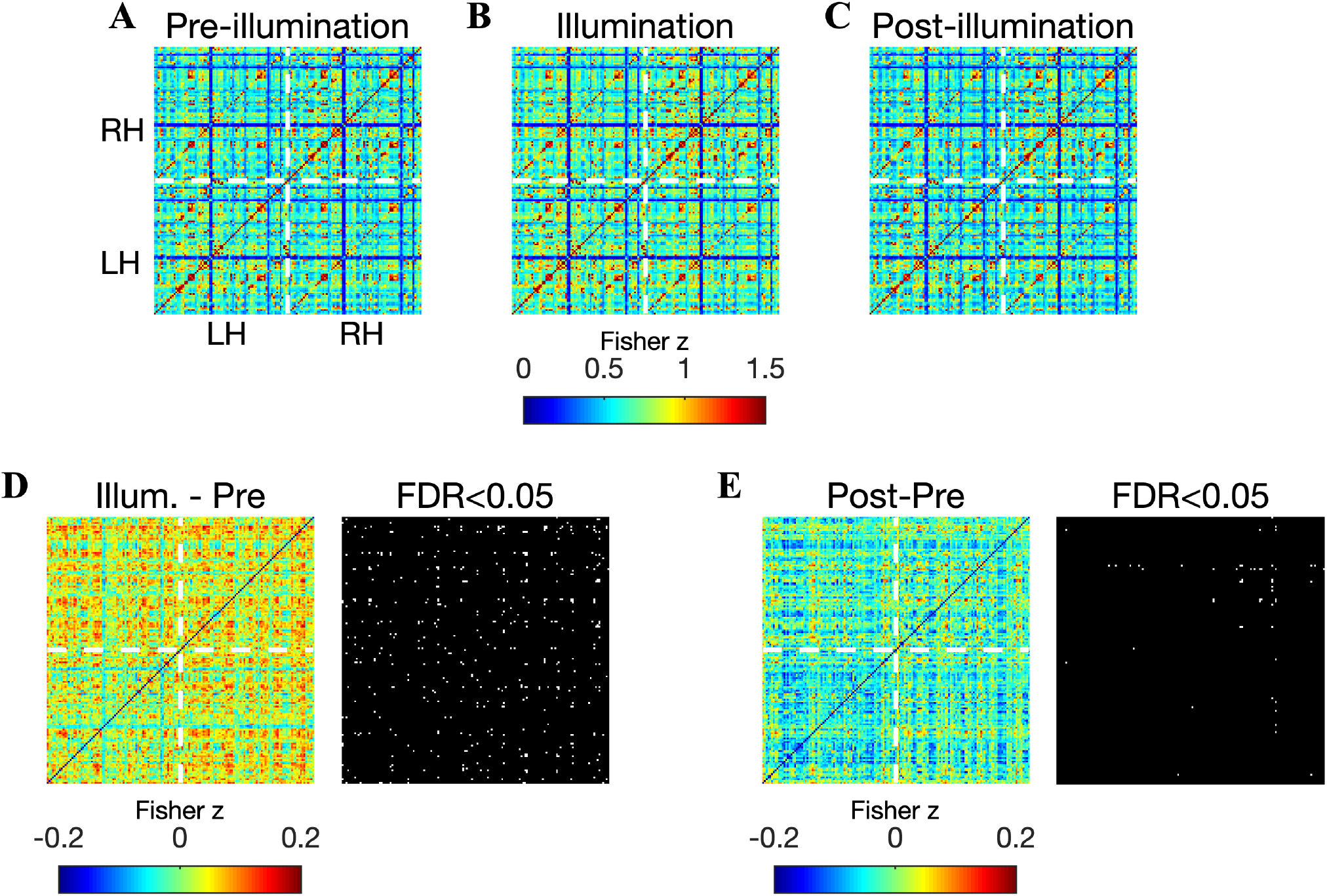
Brain-wide increases in functional connectivity during illumination. Images show the correlation matrix between all pairs of 151 ROI time courses before (**A**), during (**B**), and after (**C**) illumination. The lower left and upper right quadrants indicate connections within the left and right hemispheres, respectively. The upper left and lower right quadrants indicate interhemispheric connectivity. Stimulation was delivered to the right frontal pole. (**D**) The difference between correlation matrices measured during and before illumination: a broad increase of up to 0.2 was readily observed, with a visible dampening of the increase in the lower left quadrant – left hemispheric connections were less affected. Binary image (right) indicates the connections that exhibited a significant increase (in white) during illumination (permutation test, corrected for 11,325 comparisons using the FDR at 0.05). 219 significant connections were detected. (**E**) Same as (D) but now between the post- and pre-illumination periods. 27 connections exhibited a statistically significant increase after illumination, with a vast majority of these located within the right hemisphere.

We computed the increase in functional connectivity separately for connections within the left hemisphere, between the left and right hemispheres, and within the right hemisphere. The percent change was averaged across all connections within the specified regions: mean changes ranged from 7.3% (left hemisphere, Echo 1) to 12.8 % (interhemispheric, Echo 2), depending on the echo and hemisphere (Figure 6A-B). During illumination, we found significantly larger increases at connections involving the right hemisphere: RH-LH vs LH-LH, *p* = 0.0012 at echo 2; RH-LH vs LH-LH, *p* = 0.0359 at echo 3; RH-RH vs LH-LH, *p* = 0.030 at echo 2). The preference for the stimulated hemisphere was sustained following illumination (RH-LH vs LH-LH, *p* = 0.0026 at echo 2; RH-RH vs LH-LH *p* = 0.019 at echo 2). The number of significantly enhanced connections was also higher for connections involving the right hemisphere. During illumination at Echo 3, there 56 significantly enhanced connections within the left hemisphere, 112 between the left and right hemisphere, and 138 within the right hemisphere (Figure 6C). After illumination at echo 3, there were 2 significantly enhanced connections within the left hemisphere, 8 between the left and right hemisphere, and 34 within the right hemisphere (Figure 6D).

**Figure 6:**
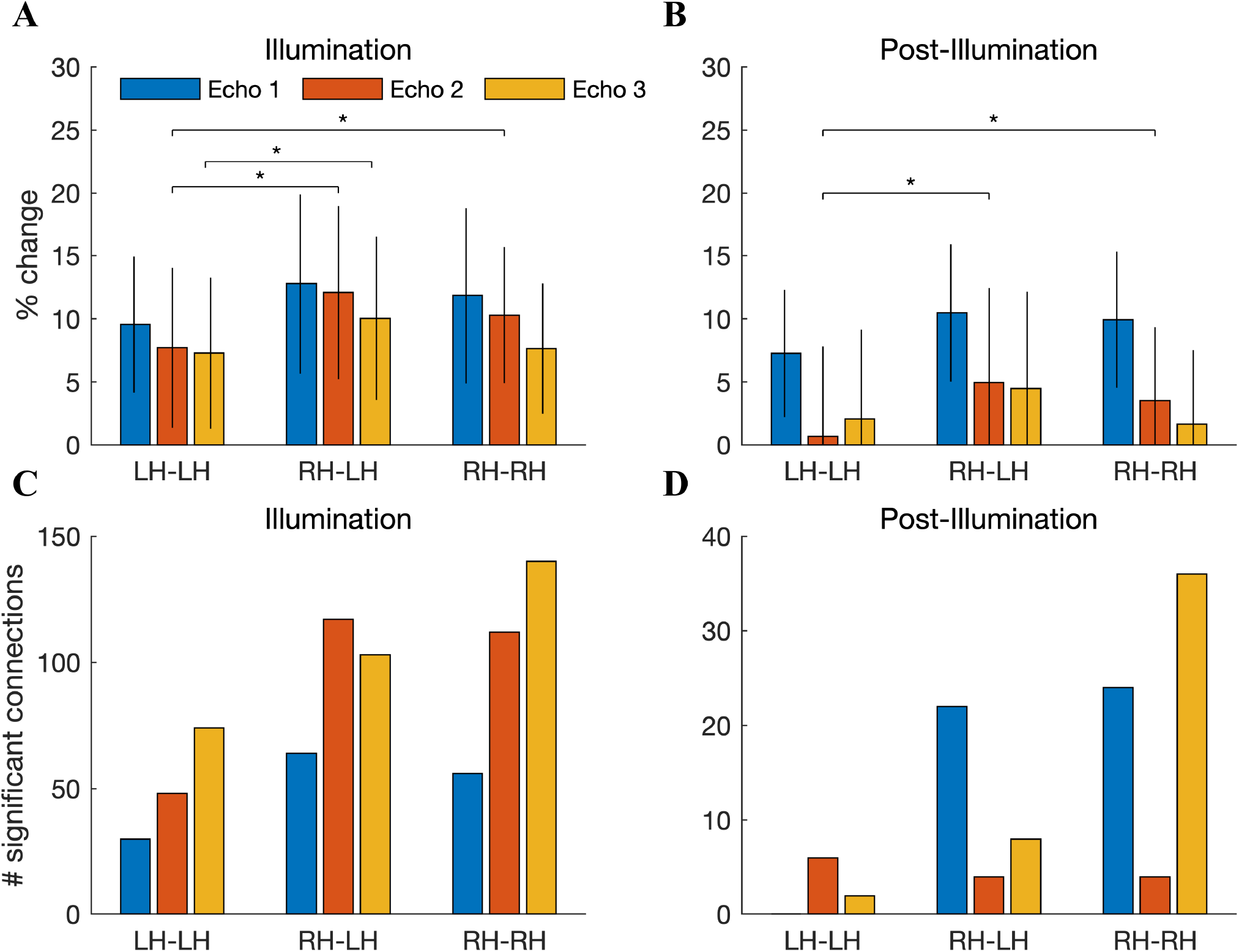
Increased functional connectivity is more pronounced in the stimulated hemisphere. (**A**) The percent increase in functional connectivity, depicted separately for connections within the left hemisphere, between the left and right hemispheres, and within the right hemisphere. The percent change was averaged across all connections in the specified region. Error bars denote the SEM across *n* = 20 subjects. During illumination, there was a significant increase in effect size for connections involving a region in the right hemisphere (RH-LH vs LH-LH: *p* = 0.0012 at echo 2; RH-LH vs LH-LH: *p* = 0.037 at echo 3; RH-RH vs LH-LH *p* = 0.032 at echo 2; paired, two-sample t-test, *n* = 20). (**B**) Connections involving the right hemisphere were also significantly enhanced after illumination (RH-LH vs LH-LH: *p* = 0.0031 at echo 2; RH-RH vs LH-LH: *p* = 0.020 at echo 2; paired, two-sample t-test). (**C**) The number of significantly enhanced connections during illumination, shown separately for left-left, left-right, and rightright connections. At all echos, the number of significantly increased connections was larger when involving a region in the right hemisphere. (**D**) Same as (C) but now for the post-illumination period. The majority of significant connections involved regions in the right hemisphere.

### No evidence for brain temperature increase with MR Thermometry

We conducted a subsequent study to test for the presence of brain temperature increases during tPBM. A separate cohort of *n* = 20 healthy subjects participated, with tPBM applied at the same dose as in the BOLD-fMRI study. Subjects performed two sessions – active and sham stimulation – in succession, with the order of active and sham randomized and balanced across subjects. We employed an MR Thermometry sequence that exploits the temperature sensitivity of the resonance frequency (Rieke and Butts Pauly, 2008; Odéen et al., 2014). A two-minute baseline period preceded the 10-minute illumination which was followed by an additional 8 minutes of recording. For both sham and active stimulation, we measured the temperature in the illuminated region at a temporal resolution of one measurement every 8.2 s (Figure 7). We did not detect significant temperature differences between active and sham stimulation at any time point before, during, or after illumination (Wilcoxon sign rank test, *n* = 20, corrected for multiple comparisons by controlling the FDR at 0.05). The maximum absolute temperature difference between active and sham stimulation was −0.14° C (i.e., a relative temperature *decrease* with active stimulation) that occurred 122s after the onset of illumination (Figure 7A). The time series of temperature in the illuminated region of individual subjects showed a high amount of overlap between active and sham stimulation (Figure 7B). We also tested for potential differences in the variability of brain temperature. The measured temperature fluctuations were very similar for active and sham stimulation (active: SD=0.14°C during stimulation; sham: SD=0.13°C; SDs averaged across *n* = 20 subjects).

**Figure 7:**
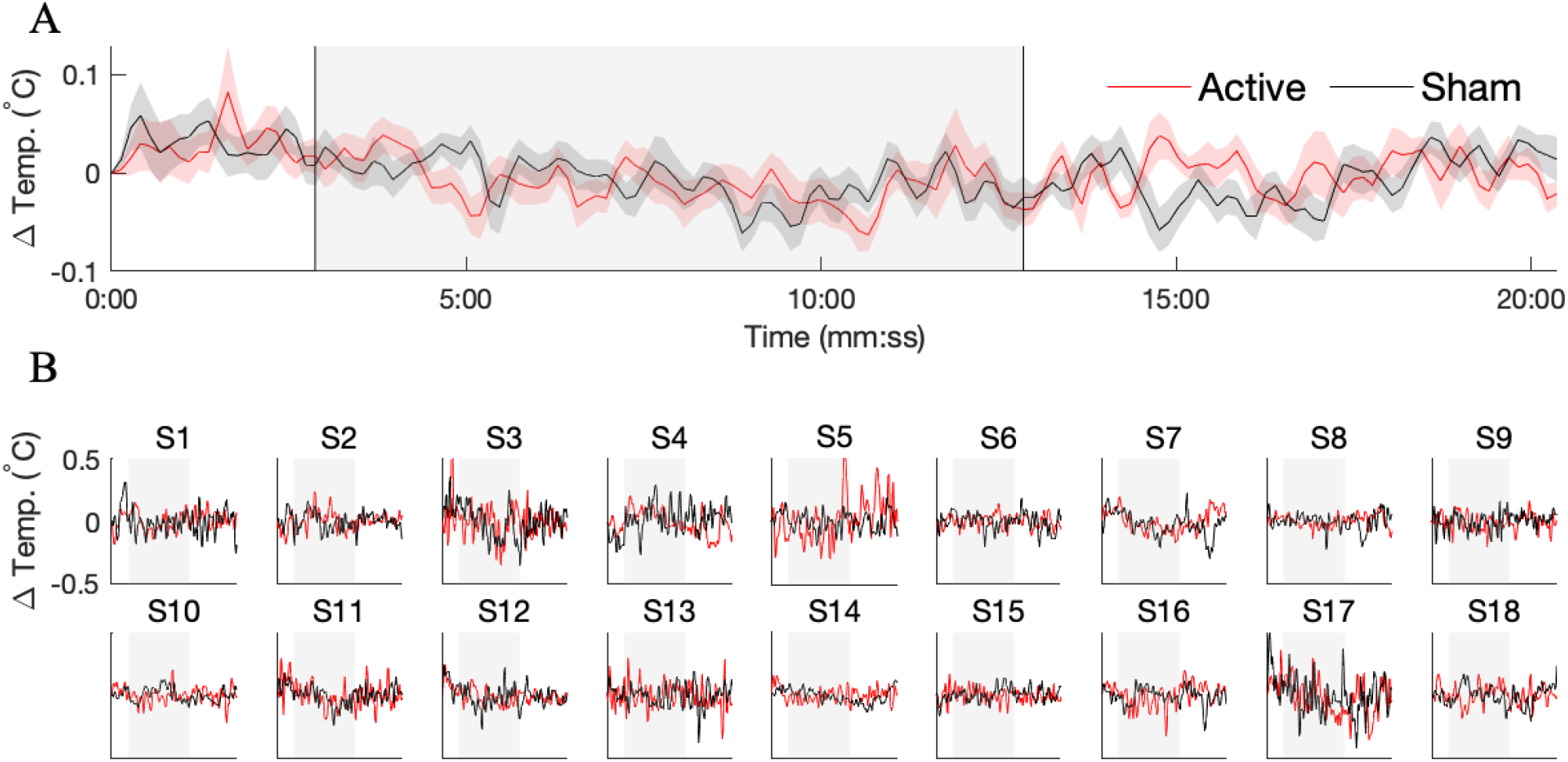
No evidence for brain heating with MR Thermometry. A separate cohort of *n* = 20 subjects was recruited for a study aimed at resolving temperature changes during tPBM with the same dose as the BOLD study. (**A**) Group-averaged temperature in the illuminated region, shown separately for active (red) and sham (black) stimulation. Error bars depict the SEM across *n* = 20 subjects. We failed to detect any time points during which the temperature in the illuminated region was significantly different during active versus sham stimulation (paired Wilcoxon signed rank test, *n* = 20, corrected for multiple comparisons by controlling the FDR at 0.05) (**B**) Temperature time courses for individual subjects. 2 of the 20 subjects were excluded due to excessive recording artifacts. Despite the presence of drifts and spurious fluctuations, the temperature time series of active and sham stimulation largely overlapped.

## Discussion

We employed BOLD-fMRI to measure hemodynamic activity in the brains of healthy human participants as they received transcranial stimulation with a near-infrared laser directed at the right frontal pole for 10 minutes. Contrary to expectation, we did not find an evoked BOLD response at illumination onset. On the other hand, we found an average 10% increase in functional connectivity with the illuminated region, indicating that hemodynamic activity in the stimulated area became more synchronized with other brain regions during illumination. The time course of connectivity exhibited a marked increase at illumination onset. Moreover, we found increased connectivity in regions outside of the directly stimulated area: functional connectivity was increased at seed regions in the frontal, temporal, and parietal cortex, with a more pronounced effect in the stimulated hemisphere. We then measured brain temperature during laser stimulation with MR Thermometry, finding no significant temperature differences between active and sham stimulation.

A limitation of our study is the difficulty in inferring the precise origin of the increased functional connectivity during illumination. The BOLD signal is a complex mixture of effects from changing CBF, CMRO2, and the arterial concentration of oxygen (Buxton, 2013). Therefore, it is challenging with the present data to disambiguate a direct vascular effect (CBF) from an indirect effect due to neural activation (CMRO2). Due to our finding of a significant BOLD effect at an early echo (13 ms), it is likely that increased CBF was involved in the observed BOLD changes – the BOLD contrast due to oxygenation is very weak at this short echo time. An inflow of fully-relaxed hemoglobin into the region would increase the initial value of the T2* signal, producing a signal increase at short echo times such as what was observed here (Posse et al., 1999; Kundu et al., 2012). On the other hand, the fact that the number of significant connections was highest at Echo 3 (i.e., 34 versus 11 in each of Echos 1 and 2, see Figure 3) suggests that cerebral oxygenation was also modulated. The change in BOLD signal due to CMRO2 increases linearly with echo time. Therefore, it seems likely that both CBF and CMRO2 were modulated during tPBM. Complicating the interpretation of a possible cascade of effects is that increased CMRO2 is expected to recruit CBF, as is the case in neural activation (Buxton, 2013).

One aspect of the present findings that points to neural activity changes during tPBM is the pattern of brain regions that was modulated. Many of the connections enhanced during illumination (e.g. posterior cingulate cortex, precuneus, inferior parietal cortex; see Tables S2–S3) belong to the “default-mode network”, a set of regions preferentially activated in the absence of a task (Van Den Heuvel and Pol, 2010; Sheline et al., 2009). A local vascular effect at the illuminated region would seem less likely to manifest in hemodynamic correlations with the set of brain areas that are believed to covary in the resting human brain. Nevertheless, to tease apart vascular and metabolic changes during tPBM, additional studies with alternative MR techniques such as arterial spin labeling (ASL) which measures CBF - see El Khoury et al. (2019) - or magnetic resonance spectroscopy which measures metabolism, are needed.

Contrary to our expectation, we failed to detect an evoked BOLD response at the onset of illumination. Due to strong low-frequency drifts in the BOLD signal, particularly when recorded over a 30-minute duration, our preprocessing included a high-pass filter that could have eliminated a very slow BOLD increase (or decrease) from the analyzed data. Another potential explanation for the absence of an evoked BOLD response is that individual response latencies varied between participants – in this case, time-locked averaging may have obscured a change in BOLD amplitude at illumination onset. The lack of an evoked response may also be related to the resting nature of the experiment: in the absence of a task-imposed metabolic need, even a hypothetical increase of mitochondrial activity may not have produced a noticeable change in the hemodynamic response. To the best of our knowledge, the downstream effect of introducing exogenous ATP into a brain region on neural signaling and vascular activity is yet unknown. Based on the robust effect on functional connectivity observed here, we speculate that the purported enhancement of mitochondrial function led to increased coordination in the activity of the stimulated region and its functionally connected regions.

Previous reports of tPBM found increases in oxygenated hemoglobin with near infrared resonance spectroscopy (NIRS) (Tian et al., 2016) and increased power of alpha band (8-12 Hz) electroencephalogram (EEG) oscillations. The large increase in functional connectivity at the latest echo time found here suggests that oxygenation was involved, consistent with (Tian et al., 2016). Moreover, associations between electrophysiological oscillations and hemodynamic activity of resting-state networks have been previously identified (Mantini et al., 2007). The present findings are distinct in the distributed nature of the hemodynamic changes observed. The fine spatial resolution afforded by fMRI allowed us to identify increased coordination among brain regions, which is difficult with EEG and NIRS, both of which are limited in spatial resolution. Thus, we have identified evidence for a “network effect” in tPBM, consistent with the fMRI changes observed in El Khoury et al. (2019).

The findings of the MR Thermometry study suggest that brain temperature does not significantly change during tPBM. The fluctuations that were observed here were within the range observed in the brains of patients following head injury, where direct measurements are feasible (Soukup et al., 2002; Wang et al., 2014). Moreover, the variability in temperature measured during tPBM (0.13-0.14 ^°^C) is similar to the temperature changes estimated from a model linking brain temperature with the BOLD signal (0.2 °C) (Yablonskiy et al., 2000). It therefore seems unlikely that the observed functional connectivity increases were mediated by a thermal effect. The absence of brain heating supports CCO as the relevant chromophore in tPBM – the most probable alternative, water, is abundant in the brain and absorption of light energy by water molecules may have led to measurable temperature changes. The PRF technique employed here to measure temperature is known to be sensitive to drift of the static magnetic field and subject motion (Rieke and Butts Pauly, 2008). We employed signal processing measures to mitigate these influences. However, it is possible that brief, acute heating followed by immediate increased CBF occurred and was not detectable within the temporal and temperature limits of our measurements. In the future, we encourage the employment of newer sequences for MR Thermometry that can achieve finer precision in temperature estimation (Odéeen and Parker, 2019). These studies will be invaluable to either conclusively rule out heating, or to identify small changes that eluded the methodology employed here.

## Acknowledgments

This research was supported by grants from the City University of New York (CUNY) and the National Aeronautics and Space Administration (NASA) to JPD. The authors would like to acknowledge Henrik Odeen and Siemens for the collaborative Work-in-progress package (WIP 1118-VE11C, Version 1.0, Nov 2017) research pulse sequence that was employed in the MR Thermometry experiments. The authors would also like to thank Lazar Fleysher for technical support during the initial development of the experimental setup. We would also like to acknowledge the help of Christian Fong in subject recruitment and experimental assistance, as well as helpful preliminary discussions with Hanli Liu.

---

**Figure S1:**
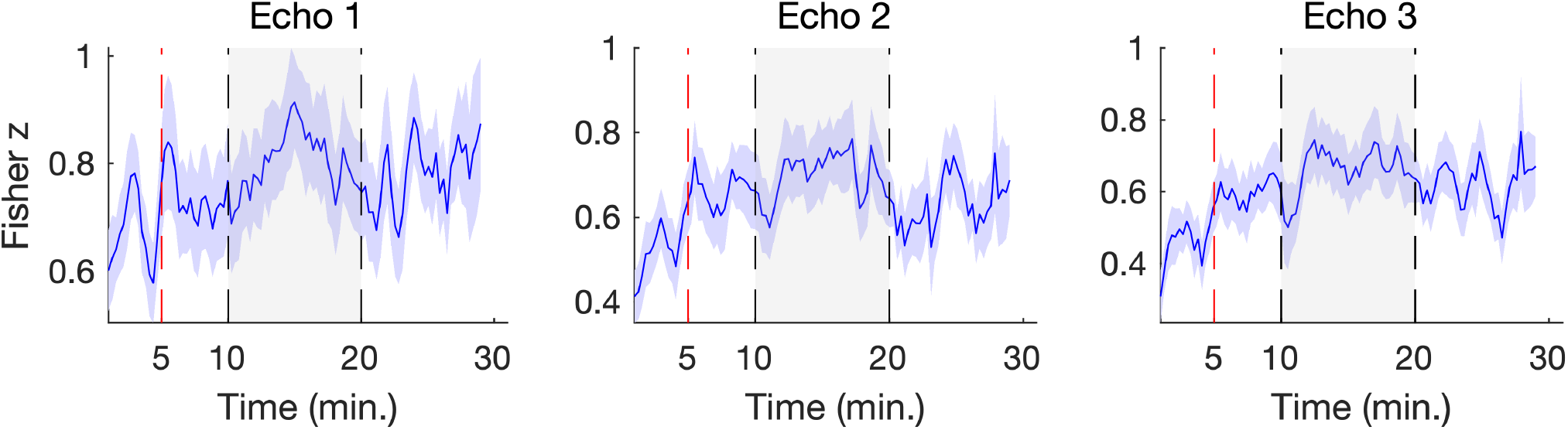
Exclusion of first 5 minutes of BOLD due to transient functional connectivity. The time course of total connectivity with the illuminated region for echo 1 (left), echo 2 (middle), and echo 3 (right). Connectivity exhibited transient oscillations during the first five minutes of the pre-illumination period before reaching a stable value at approximately five minutes (indicated with vertical dashed red line). Consequently, we excluded the first five minutes of the baseline period from further analysis.

**Figure S2:**
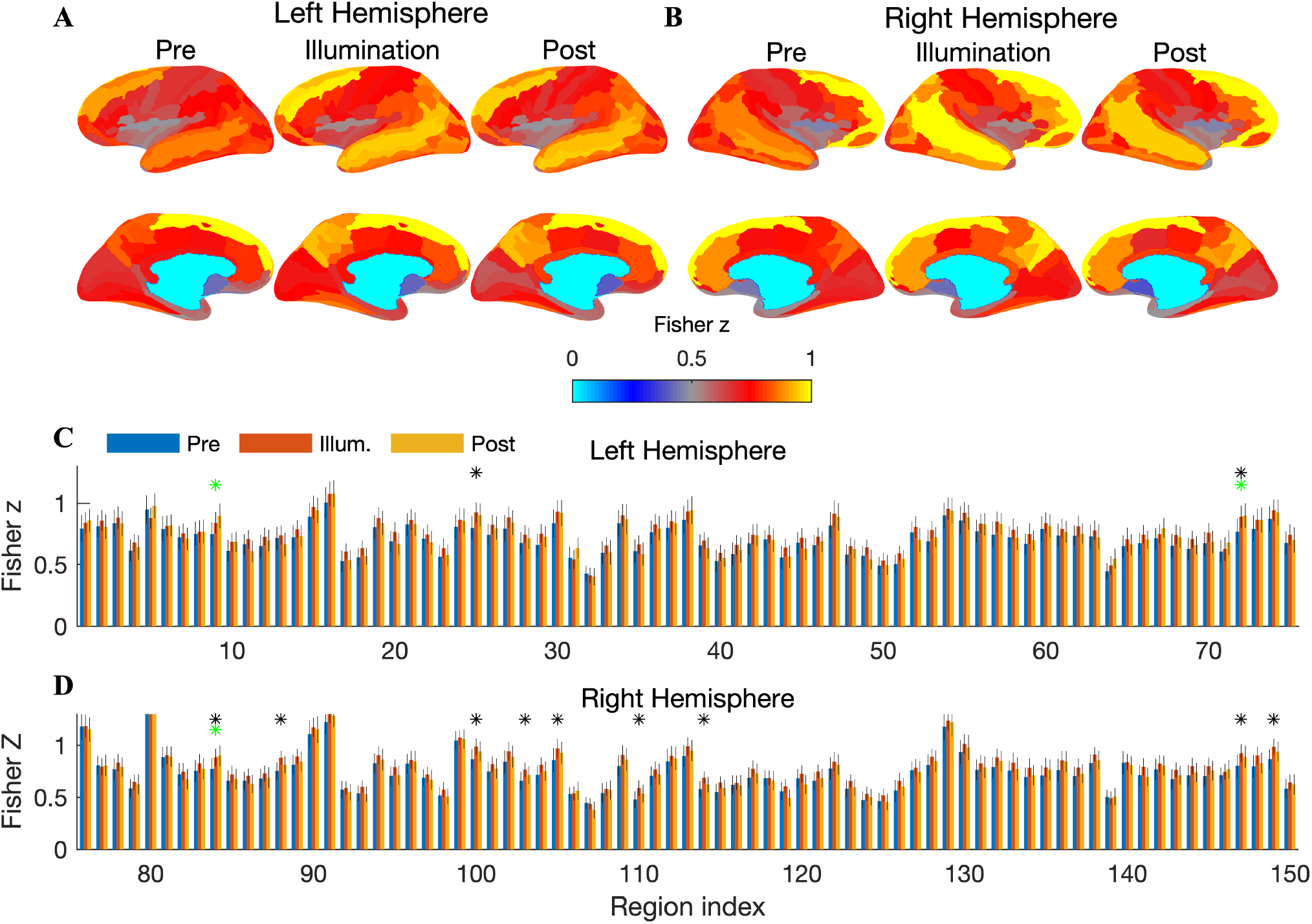
Increased functional connectivity with the illuminated region at Echo 2. (**A**) Cortical surfaces depict the Fisher transformed Pearson correlation between the illuminated region and all ROIs in the left hemisphere, shown separately for the pre-illumination, illumination, and post-illumination period. Increased connectivity over the frontal, temporal, and parietal cortex is apparent. (**B**) Same as (A) but now showing connectivity between the illuminated region and ROIs in the right hemisphere. (**C**) Bar graphs depict the mean connectivity before, during, and after illumination for each ROI in the left hemisphere. Asterisks denote a significant increase during illumination. (**D**) Same as (C) but now showing connectivity with the right hemisphere. In total, 73 ROIs exhibited a significant increase in connectivity with the illuminated region.

**Figure S3:**
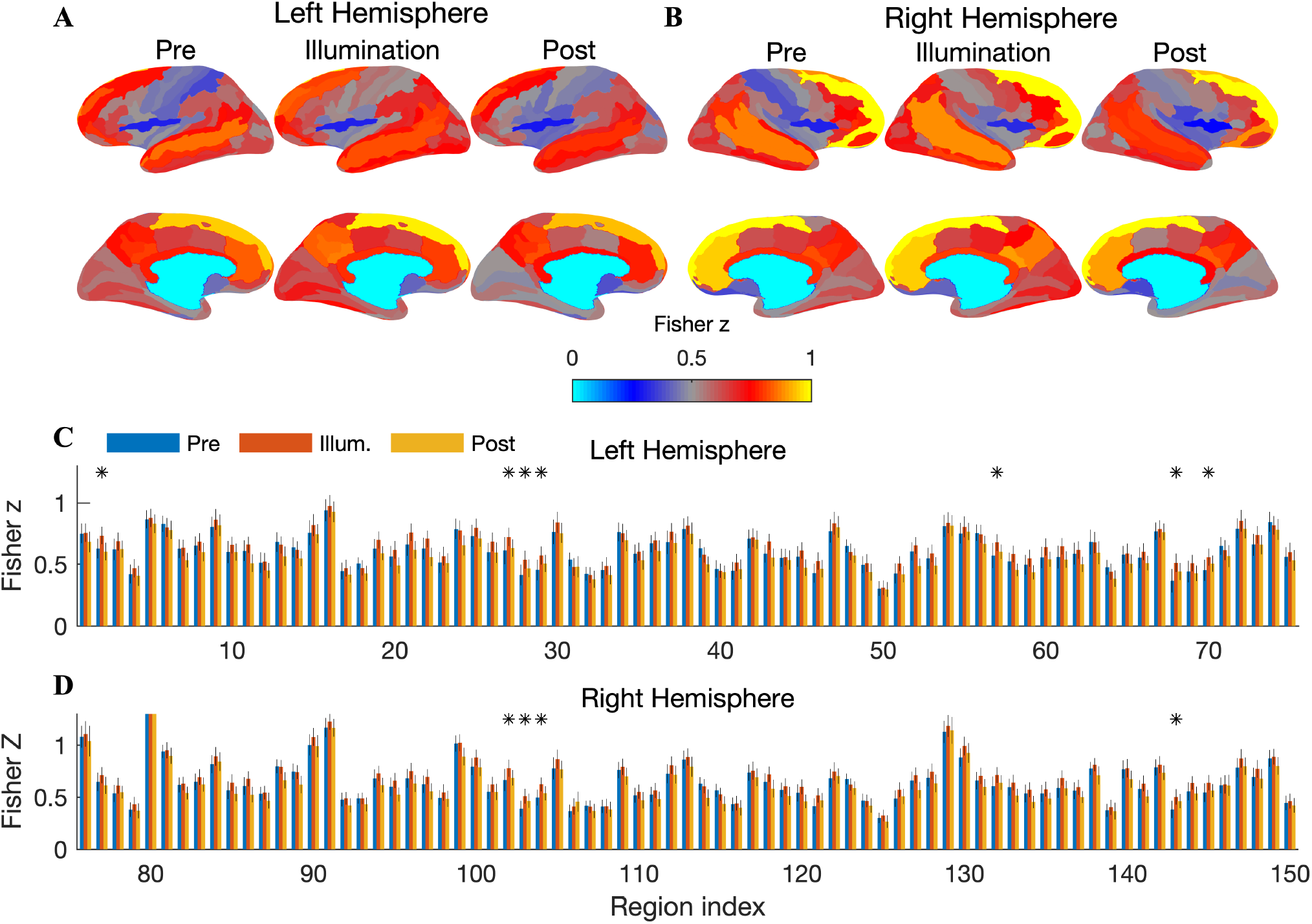
Increased functional connectivity with the illuminated region at Echo 3. (**A**) Connections with the the frontal, temporal, and parietal cortex exhibit stronger correlations. (**B**) Same as (A) but now showing connectivity between the illuminated region and ROIs in the right hemisphere. (**C**) The mean connectivity before, during, and after illumination, shown separately for each ROI in the left hemisphere. Asterisks denote a significant increase during illumination. (**D**) Same as (C) but now showing connectivity with the right hemisphere. In total, 87 ROIs exhibited a significant increase in connectivity with the illuminated region.

**Table S1:** Physical and optical properties of the four-layer head model employed in the simulation of light propagation. Whenever possible, we have used *in vivo* optical properties. In particular, while MCML requires separate specification of *μ_s_* and *g*, *in vivo* values for the reduced scattering coefficient 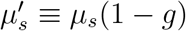 and *g* are more readily available. As such, we have inferred values of *μ_s_* from reported values of 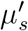 and *g*. The references cited for *μ_s_* report *in vivo* values of reduced scattering coefficients. All values of *μ_a_* and *μ_s_* are in units of cm^-1^. A grid spacing of 20*μ*m was used in both radial and axial directions, and 500,000 photons were simulated in each of five runs. We report the average absorption across the 5 runs.

| Layer | Refractive Index | Absorption Coefficient ($\mu_a$) | Scattering Coefficient ($\mu_s$) | Anisotropy Factor ($g$) | Thickness (cm) |
| --- | --- | --- | --- | --- | --- |
| Scalp | 1.41 | 0.2 | 31.25 | 0.52 | 0.6 |
| Skull | 1.56 | 0.07 | 130 | 0.9 | 0.65 |
| CSF | 1.33 | 0.02 | 0.1 | 0.9 | 0.2 |
| Brain | 1.4 | 0.084 | 90 | 0.9 | 8 |

**Figure S4:**
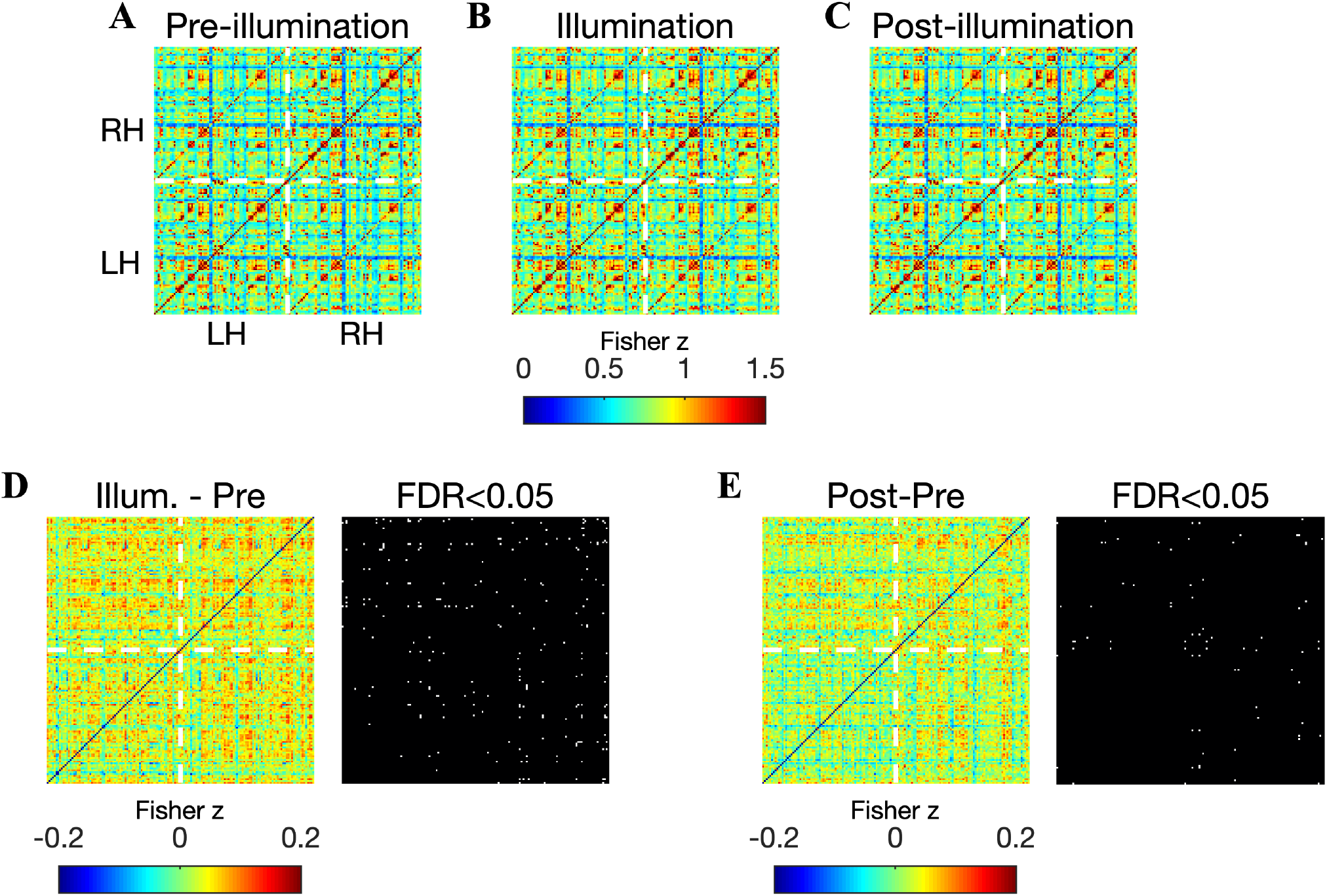
Increased functional connectivity with seeds outside of the illuminated region. (**A**) Matrix showing the Pearson correlation between all pairs of 151 ROI time courses before illumination (echo 2). (**B**) Same as (A) but now for the illumination period. Increased correlations are apparent throughout broad sections of the matrix. (**C**) Correlation matrix for the period after illumination, where the increased correlation is dampened. (**D**) The difference in correlation matrices measured during and pre-illumination: a broad increase of up to 0.2 is readily observed. (**E**) Same as (D) but now between the post- and pre-illumination periods. Only sparse increases are apparent. (**F**) The total connectivity was computed for all seeds and then sorted in descending order of correlation difference between illumination and pre-illumination. The 5 seeds showing the largest increase in functional connectivity were all in the frontal cortex (color-coded, see legend). Notably, the illuminated region had the fourth strongest effect out of 151 seeds. 67 of the 151 seeds exhibited significantly increased total connectivity during the illumination period (indicated with blue bars; paired t-test on Fisher transformed correlation coefficients, *n* = 20, corrected for multiple comparisons by controlling the FDR at 0.05). At echos 1 and 3, functional connectivity increases at seeds outside of the illuminated region did not pass statistical significance after correcting for multiple comparisons.

**Figure S5:**
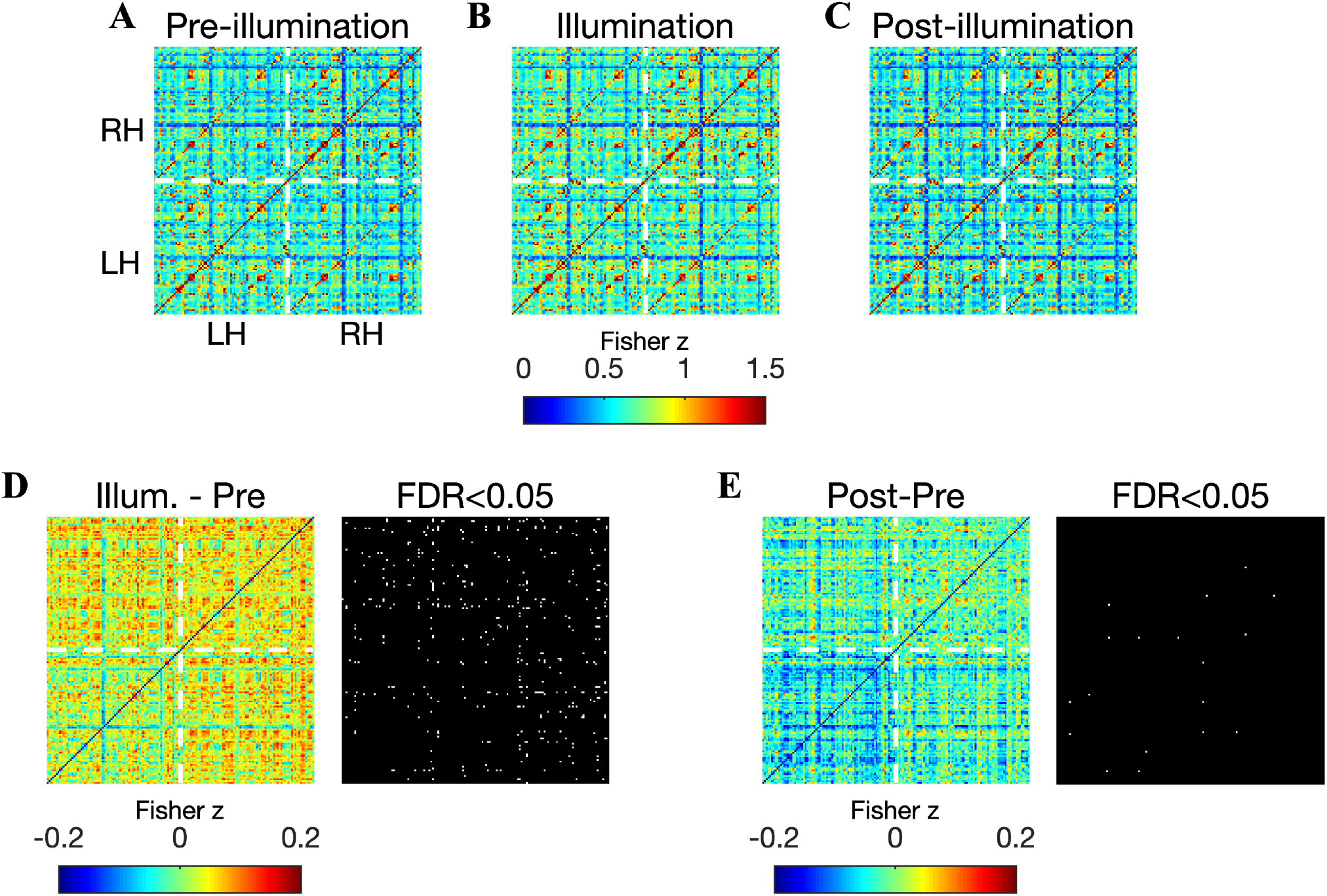
Increased functional connectivity with seeds outside of the illuminated region. (**A**) Matrix showing the Pearson correlation between all pairs of 151 ROI time courses before illumination (echo 2). (**B**) Same as (A) but now for the illumination period. Increased correlations are apparent throughout broad sections of the matrix. (**C**) Correlation matrix for the period after illumination, where the increased correlation is dampened. (**D**) The difference in correlation matrices measured during and pre-illumination: a broad increase of up to 0.2 is readily observed. (**E**) Same as (D) but now between the post- and pre-illumination periods. Only sparse increases are apparent. (**F**) The total connectivity was computed for all seeds and then sorted in descending order of correlation difference between illumination and pre-illumination. The 5 seeds showing the largest increase in functional connectivity were all in the frontal cortex (color-coded, see legend). Notably, the illuminated region had the fourth strongest effect out of 151 seeds. 67 of the 151 seeds exhibited significantly increased total connectivity during the illumination period (indicated with blue bars; paired t-test on Fisher transformed correlation coefficients, *n* = 20, corrected for multiple comparisons by controlling the FDR at 0.05). At echos 1 and 3, functional connectivity increases at seeds outside of the illuminated region did not pass statistical significance after correcting for multiple comparisons.

**Table S2:** Functional connectivity between the illuminated region and all connecting ROIs in the left hemisphere. Connections exhibiting a statistically significant increase during or after illumination are indicated with bold font. There were 2, 6, and 21 significantly enhanced connections during illuminations in echos 1, 2, and 3, respectively (permutation test, corrected for multiple comparisons by controlling the FDR at 0.05.)

| Connection | Pre | Echo 1<br>Illum. | Post | Pre | Echo 2<br>Illum. | Post | Pre | Echo 3<br>Illum. | Post |
| --- | --- | --- | --- | --- | --- | --- | --- | --- | --- |
| G&S.frontomargin | 0.80±0.11 | 0.84±0.09 | 0.86±0.09 | 0.75±0.08 | 0.76±0.08 | 0.68±0.09 | 0.61±0.08 | **0.72±0.07** | 0.70±0.08 |
| G&S.occipital_inf | 0.82±0.09 | 0.86±0.08 | 0.81±0.10 | 0.63±0.08 | **0.74±0.07** | 0.60±0.07 | 0.55±0.07 | **0.68±0.07** | 0.62±0.07 |
| G&S.paracentral | 0.84±0.11 | 0.88±0.09 | 0.84±0.10 | 0.62±0.09 | 0.69±0.08 | 0.62±0.07 | 0.58±0.07 | 0.68±0.08 | 0.65±0.07 |
| G&S.subcentral | 0.61±0.09 | 0.68±0.08 | 0.64±0.10 | 0.42±0.08 | 0.47±0.07 | 0.41±0.08 | 0.40±0.07 | 0.46±0.08 | 0.46±0.08 |
| G&S.transv_frontopol | 0.95±0.11 | 0.88±0.09 | 0.98±0.10 | 0.87±0.07 | 0.88±0.07 | 0.83±0.08 | 0.67±0.08 | 0.77±0.07 | 0.78±0.08 |
| G&S.cingul-Ant | 0.79±0.10 | 0.82±0.08 | 0.82±0.09 | 0.83±0.07 | 0.80±0.08 | 0.78±0.08 | 0.71±0.06 | 0.76±0.08 | 0.74±0.09 |
| G&S.cingul-Mid-Ant | 0.72±0.10 | 0.76±0.08 | 0.71±0.09 | 0.62±0.08 | 0.63±0.07 | 0.53±0.06 | 0.55±0.07 | 0.60±0.08 | 0.60±0.05 |
| G&S.cingul-Mid-Post | 0.75±0.10 | 0.77±0.10 | 0.77±0.10 | 0.65±0.08 | 0.68±0.07 | 0.60±0.07 | 0.60±0.07 | 0.65±0.08 | 0.64±0.07 |
| G.cingul-Post-dorsal | 0.75±0.10 | 0.84±0.09 | **0.90±0.10** | 0.81±0.08 | 0.87±0.09 | 0.82±0.09 | 0.70±0.08 | 0.81±0.09 | 0.81±0.09 |
| G.cingul-Post-ventral | 0.61±0.09 | 0.68±0.08 | 0.68±0.09 | 0.60±0.09 | 0.66±0.07 | 0.60±0.07 | 0.50±0.09 | **0.65±0.07** | 0.61±0.07 |
| G.cuneus | 0.66±0.09 | 0.71±0.08 | 0.61±0.09 | 0.61±0.08 | 0.66±0.07 | 0.51±0.07 | 0.52±0.08 | **0.64±0.07** | 0.54±0.07 |
| G.front_inf-Opercular | 0.65±0.10 | 0.73±0.07 | 0.70±0.09 | 0.51±0.08 | 0.52±0.07 | 0.45±0.06 | 0.52±0.08 | 0.52±0.07 | 0.50±0.06 |
| G.front_inf-Orbital | 0.71±0.10 | 0.74±0.08 | 0.66±0.10 | 0.68±0.08 | 0.66±0.07 | 0.56±0.08 | 0.60±0.09 | 0.63±0.08 | 0.61±0.08 |
| G.front_inf-Triangul | 0.72±0.10 | 0.79±0.08 | 0.74±0.09 | 0.64±0.08 | 0.62±0.07 | 0.55±0.06 | 0.60±0.08 | 0.61±0.07 | 0.59±0.06 |
| G.front_middle | 0.89±0.11 | 0.97±0.09 | 0.94±0.10 | 0.76±0.10 | 0.82±0.09 | 0.75±0.08 | 0.70±0.09 | 0.79±0.09 | 0.75±0.08 |
| G.front_sup | 1.01±0.13 | 1.08±0.10 | 1.08±0.11 | 0.94±0.09 | 0.98±0.09 | 0.93±0.08 | 0.86±0.09 | 0.91±0.09 | 0.93±0.09 |
| G.Ins.lg&S.cent.ins | 0.53±0.09 | 0.60±0.06 | 0.54±0.08 | 0.44±0.07 | 0.47±0.07 | 0.42±0.07 | 0.45±0.06 | 0.51±0.07 | 0.48±0.07 |
| G.insular_short | 0.56±0.08 | 0.63±0.06 | 0.57±0.07 | 0.51±0.05 | 0.47±0.06 | 0.43±0.06 | 0.52±0.05 | 0.49±0.07 | 0.48±0.07 |
| G.occipital_middle | 0.81±0.10 | 0.88±0.09 | 0.84±0.10 | 0.63±0.08 | 0.70±0.07 | 0.59±0.06 | 0.54±0.07 | **0.66±0.08** | 0.61±0.06 |
| G.occipital_sup | 0.69±0.09 | 0.77±0.08 | 0.67±0.09 | 0.56±0.09 | 0.62±0.07 | 0.49±0.07 | 0.49±0.07 | 0.60±0.07 | 0.50±0.06 |
| G.oc-temp.lat-fusiform | 0.83±0.09 | 0.87±0.09 | 0.83±0.10 | 0.66±0.09 | 0.76±0.07 | 0.62±0.07 | 0.57±0.08 | **0.70±0.08** | 0.62±0.07 |
| G.oc-temp.med-Lingual | 0.71±0.09 | 0.75±0.08 | 0.68±0.10 | 0.63±0.09 | 0.71±0.07 | 0.56±0.08 | 0.56±0.08 | **0.69±0.07** | 0.59±0.07 |
| G.oc-temp.med-Parahip | 0.56±0.09 | 0.63±0.08 | 0.58±0.08 | 0.51±0.09 | 0.56±0.08 | 0.51±0.08 | 0.47±0.07 | 0.52±0.07 | 0.46±0.07 |
| G.orbital | 0.81±0.10 | 0.87±0.10 | 0.86±0.10 | 0.79±0.09 | 0.78±0.08 | 0.65±0.08 | 0.65±0.08 | 0.69±0.08 | 0.64±0.08 |
| G.pariet_inf-Angular | 0.80±0.10 | **0.93±0.08** | 0.91±0.09 | 0.73±0.09 | 0.80±0.07 | 0.71±0.07 | 0.67±0.09 | 0.77±0.08 | 0.70±0.07 |
| G.pariet_inf-Supramar | 0.75±0.09 | 0.83±0.08 | 0.80±0.10 | 0.60±0.07 | 0.68±0.08 | 0.59±0.08 | 0.57±0.07 | 0.65±0.09 | 0.63±0.08 |
| G.parietal_sup | 0.80±0.11 | 0.89±0.09 | 0.84±0.10 | 0.61±0.11 | **0.72±0.08** | 0.63±0.07 | 0.55±0.10 | **0.69±0.08** | 0.63±0.07 |
| G.postcentral | 0.67±0.09 | 0.75±0.08 | 0.71±0.09 | 0.41±0.08 | **0.54±0.07** | 0.47±0.07 | 0.41±0.07 | **0.53±0.08** | 0.52±0.07 |
| G.precentral | 0.66±0.10 | 0.75±0.08 | 0.73±0.09 | 0.45±0.08 | **0.57±0.07** | 0.50±0.07 | 0.43±0.07 | **0.55±0.08** | 0.55±0.07 |
| G.precuneus | 0.84±0.11 | 0.93±0.09 | 0.92±0.11 | 0.76±0.10 | 0.84±0.09 | 0.76±0.09 | 0.67±0.09 | **0.80±0.09** | 0.77±0.09 |
| G.rectus | 0.55±0.09 | 0.54±0.08 | 0.63±0.09 | 0.54±0.06 | 0.48±0.07 | 0.48±0.09 | 0.27±0.06 | 0.27±0.07 | 0.34±0.08 |
| G.subcallosal | 0.43±0.06 | 0.41±0.06 | 0.40±0.07 | 0.42±0.06 | 0.41±0.06 | 0.38±0.07 | 0.25±0.05 | 0.21±0.05 | 0.22±0.06 |
| G.temp_sup-G.T.transv | 0.59±0.09 | 0.65±0.07 | 0.60±0.10 | 0.45±0.07 | 0.49±0.07 | 0.41±0.08 | 0.42±0.06 | 0.50±0.08 | 0.48±0.07 |
| G.temp_sup-Lateral | 0.84±0.11 | 0.90±0.09 | 0.87±0.12 | 0.76±0.08 | 0.76±0.08 | 0.69±0.09 | 0.70±0.08 | 0.71±0.08 | 0.69±0.09 |
| G.temp_sup-Plan.polar | 0.61±0.10 | 0.66±0.08 | 0.58±0.09 | 0.59±0.07 | 0.60±0.07 | 0.53±0.08 | 0.55±0.07 | 0.55±0.07 | 0.50±0.09 |
| G.temp_sup-Plan.temporo | 0.76±0.10 | 0.83±0.09 | 0.80±0.10 | 0.67±0.07 | 0.69±0.08 | 0.61±0.09 | 0.62±0.07 | 0.66±0.08 | 0.63±0.09 |
| G.temporal_inf | 0.80±0.11 | 0.85±0.09 | 0.84±0.10 | 0.68±0.10 | 0.77±0.07 | 0.67±0.08 | 0.55±0.08 | **0.66±0.08** | 0.61±0.08 |
| G.temporal_middle | 0.87±0.11 | 0.94±0.10 | 0.94±0.11 | 0.79±0.09 | 0.82±0.07 | 0.75±0.09 | 0.68±0.07 | 0.74±0.08 | 0.70±0.09 |
| Lat.Fis-ant-Horizont | 0.65±0.09 | 0.69±0.07 | 0.63±0.09 | 0.63±0.07 | 0.58±0.06 | 0.50±0.07 | 0.56±0.07 | 0.57±0.07 | 0.57±0.07 |
| Lat.Fis-ant-Vertical | 0.53±0.09 | 0.59±0.07 | 0.55±0.07 | 0.46±0.07 | 0.45±0.06 | 0.43±0.06 | 0.49±0.06 | 0.48±0.06 | 0.50±0.05 |
| Lat.Fis-post | 0.58±0.09 | 0.66±0.08 | 0.62±0.10 | 0.45±0.07 | 0.51±0.07 | 0.46±0.08 | 0.44±0.07 | 0.52±0.08 | 0.51±0.08 |
| Medial.wall | 0.67±0.09 | 0.75±0.08 | 0.74±0.09 | 0.71±0.08 | 0.72±0.08 | 0.70±0.08 | 0.58±0.08 | 0.62±0.08 | 0.58±0.08 |
| Pole.occipital | 0.70±0.08 | 0.74±0.07 | 0.70±0.10 | 0.59±0.07 | 0.68±0.07 | 0.55±0.07 | 0.54±0.07 | **0.67±0.07** | 0.58±0.07 |
| Pole.temporal | 0.56±0.10 | 0.64±0.07 | 0.56±0.09 | 0.55±0.07 | 0.56±0.07 | 0.53±0.08 | 0.50±0.07 | 0.49±0.06 | 0.42±0.06 |
| S.calcarine | 0.68±0.09 | 0.72±0.08 | 0.62±0.09 | 0.56±0.08 | 0.61±0.07 | 0.47±0.07 | 0.48±0.07 | 0.60±0.07 | 0.51±0.06 |
| S.central | 0.66±0.09 | 0.73±0.07 | 0.69±0.09 | 0.43±0.08 | 0.52±0.07 | 0.46±0.07 | 0.41±0.06 | **0.51±0.08** | 0.51±0.07 |
| S.cingul-Marginalis | 0.82±0.11 | 0.92±0.11 | 0.89±0.11 | 0.77±0.10 | 0.84±0.09 | 0.80±0.09 | 0.73±0.09 | 0.82±0.10 | 0.82±0.10 |
| S.circular.insula_ant | 0.58±0.08 | 0.65±0.07 | 0.62±0.08 | 0.65±0.05 | 0.60±0.07 | 0.57±0.06 | 0.61±0.07 | 0.59±0.08 | 0.57±0.08 |
| S.circular.insula_inf | 0.57±0.08 | 0.64±0.06 | 0.54±0.08 | 0.50±0.07 | 0.51±0.07 | 0.44±0.07 | 0.49±0.07 | 0.55±0.08 | 0.52±0.08 |
| S.circular.insula_sup | 0.49±0.07 | 0.53±0.06 | 0.50±0.08 | 0.30±0.06 | 0.31±0.06 | 0.29±0.06 | 0.36±0.06 | 0.36±0.07 | 0.37±0.06 |
| S.collat.transv_ant | 0.50±0.08 | 0.59±0.06 | 0.54±0.07 | 0.42±0.08 | 0.50±0.06 | 0.42±0.07 | 0.36±0.06 | 0.41±0.06 | 0.34±0.05 |
| S.collat.transv_post | 0.76±0.09 | 0.81±0.09 | 0.70±0.10 | 0.60±0.08 | 0.65±0.07 | 0.49±0.06 | 0.54±0.07 | 0.63±0.07 | 0.51±0.07 |
| S.front_inf | 0.69±0.10 | 0.78±0.07 | 0.72±0.08 | 0.55±0.09 | 0.59±0.07 | 0.49±0.07 | 0.52±0.08 | 0.59±0.07 | 0.53±0.07 |
| S.front_middle | 0.90±0.11 | 0.96±0.09 | 0.94±0.11 | 0.81±0.09 | 0.84±0.09 | 0.82±0.09 | 0.75±0.09 | 0.83±0.09 | 0.84±0.08 |
| S.front_sup | 0.86±0.11 | 0.92±0.09 | 0.89±0.10 | 0.75±0.09 | 0.81±0.09 | 0.77±0.07 | 0.69±0.09 | 0.77±0.10 | 0.78±0.08 |
| S.interm_prim-Jensen | 0.77±0.10 | 0.84±0.08 | 0.83±0.09 | 0.76±0.07 | 0.75±0.08 | 0.66±0.07 | 0.72±0.08 | 0.74±0.09 | 0.66±0.07 |
| S.intrapariet&P.trans | 0.75±0.11 | 0.85±0.08 | 0.83±0.09 | 0.57±0.10 | **0.68±0.07** | 0.60±0.06 | 0.53±0.09 | **0.65±0.07** | 0.60±0.06 |
| S.oc.middle&Lunatus | 0.72±0.09 | 0.79±0.09 | 0.72±0.10 | 0.52±0.07 | 0.58±0.07 | 0.45±0.05 | 0.47±0.06 | 0.56±0.07 | 0.49±0.05 |
| S.oc_sup&transversal | 0.67±0.09 | 0.75±0.09 | 0.70±0.09 | 0.50±0.08 | 0.55±0.07 | 0.43±0.06 | 0.44±0.07 | 0.53±0.07 | 0.46±0.05 |
| S.occipital_ant | 0.79±0.10 | 0.84±0.09 | 0.82±0.10 | 0.56±0.09 | 0.64±0.07 | 0.54±0.06 | 0.49±0.08 | 0.61±0.07 | 0.56±0.06 |
| S.oc-temp.lat | 0.74±0.10 | 0.79±0.08 | 0.77±0.09 | 0.56±0.09 | 0.64±0.07 | 0.54±0.07 | 0.47±0.08 | 0.58±0.07 | 0.50±0.06 |
| S.oc-temp.med&Lingual | 0.74±0.09 | 0.81±0.09 | 0.75±0.08 | 0.59±0.09 | 0.62±0.07 | 0.50±0.06 | 0.53±0.07 | 0.60±0.07 | 0.52±0.06 |
| S.orbital.lateral | 0.73±0.10 | 0.78±0.08 | 0.73±0.10 | 0.68±0.10 | 0.68±0.09 | 0.59±0.09 | 0.61±0.10 | 0.68±0.09 | 0.64±0.09 |
| S.orbital.med-olfact | 0.45±0.07 | 0.49±0.08 | 0.55±0.08 | 0.47±0.06 | 0.44±0.07 | 0.38±0.07 | 0.22±0.05 | 0.18±0.06 | 0.24±0.08 |
| S.orbital-H.Shaped | 0.65±0.09 | 0.70±0.10 | 0.66±0.09 | 0.58±0.07 | 0.59±0.08 | 0.50±0.07 | 0.46±0.06 | 0.48±0.07 | 0.41±0.08 |
| S.parieto-occipital | 0.67±0.09 | 0.75±0.08 | 0.70±0.08 | 0.55±0.09 | 0.60±0.07 | 0.51±0.06 | 0.49±0.08 | 0.60±0.07 | 0.53±0.06 |
| S.pericallosal | 0.71±0.09 | 0.75±0.08 | 0.80±0.08 | 0.77±0.07 | 0.79±0.07 | 0.76±0.07 | 0.68±0.08 | 0.76±0.08 | 0.74±0.08 |
| S.postcentral | 0.65±0.10 | 0.74±0.08 | 0.72±0.09 | 0.37±0.09 | **0.51±0.08** | 0.44±0.07 | 0.38±0.08 | **0.50±0.08** | 0.48±0.06 |
| S.precentral-inf-part | 0.62±0.09 | 0.71±0.07 | 0.66±0.08 | 0.44±0.09 | 0.51±0.07 | 0.43±0.06 | 0.43±0.08 | 0.49±0.07 | 0.46±0.07 |
| S.precentral-sup-part | 0.67±0.10 | 0.76±0.08 | 0.68±0.09 | 0.45±0.09 | **0.56±0.07** | 0.50±0.07 | 0.41±0.07 | **0.52±0.08** | **0.53±0.07** |
| S.suborbital | 0.60±0.09 | 0.63±0.10 | 0.68±0.10 | 0.65±0.07 | 0.61±0.08 | 0.57±0.09 | 0.38±0.06 | 0.42±0.08 | 0.43±0.09 |
| S.subparietal | 0.77±0.11 | **0.89±0.09** | **0.90±0.10** | 0.79±0.08 | 0.86±0.09 | 0.79±0.08 | 0.72±0.09 | 0.83±0.09 | 0.81±0.09 |
| S.temporal_inf | 0.79±0.11 | 0.87±0.09 | 0.87±0.10 | 0.66±0.09 | 0.74±0.07 | 0.66±0.08 | 0.53±0.07 | 0.62±0.07 | 0.57±0.07 |
| S.temporal_sup | 0.87±0.10 | 0.94±0.09 | 0.93±0.11 | 0.85±0.08 | 0.82±0.08 | 0.78±0.09 | 0.74±0.08 | 0.78±0.08 | 0.77±0.09 |
| S.temporal_transverse | 0.67±0.10 | 0.75±0.09 | 0.70±0.11 | 0.56±0.08 | 0.60±0.08 | 0.53±0.08 | 0.50±0.07 | 0.57±0.08 | 0.56±0.08 |

**Table S3:** Functional connectivity between the illuminated region and all connecting ROIs in the right hemisphere. 6, 6, and 417 significantly enhanced connections were found at echos 1, 2, and 3, respectively, during illumination (permutation test, corrected for multiple comparisons by controlling the FDR at 0.05.)

| Connection | Echo 1 |  |  | Echo 2 |  |  | Echo 3 |  |  |
| --- | --- | --- | --- | --- | --- | --- | --- | --- | --- |
|  | Pre | Illum. | Post | Pre | Illum. | Post | Pre | Illum. | Post |
| G&S.frontomargin | 1.18±0.13 | 1.18±0.11 | 1.15±0.12 | 1.08±0.10 | 1.11±0.12 | 1.04±0.15 | 1.06±0.11 | 1.04±0.10 | 1.02±0.12 |
| G&S.occipital_inf | 0.81±0.09 | 0.79±0.08 | 0.81±0.10 | 0.65±0.09 | 0.71±0.08 | 0.61±0.08 | 0.58±0.08 | 0.67±0.08 | 0.62±0.07 |
| G&S.paracentral | 0.77±0.10 | 0.83±0.08 | 0.79±0.09 | 0.54±0.08 | 0.61±0.08 | 0.55±0.06 | 0.49±0.07 | 0.60±0.08 | 0.57±0.07 |
| G&S.subcentral | 0.59±0.09 | 0.65±0.07 | 0.62±0.10 | 0.38±0.07 | 0.43±0.07 | 0.37±0.07 | 0.39±0.06 | 0.43±0.08 | 0.42±0.08 |
| G&S.transv.frontopol | 1.53±0.11 | 1.52±0.11 | 1.56±0.13 | 1.55±0.12 | 1.56±0.12 | 1.50±0.14 | 1.50±0.13 | 1.54±0.13 | 1.52±0.14 |
| G&S.cingul-Ant | 0.89±0.10 | 0.90±0.09 | 0.89±0.10 | 0.94±0.06 | 0.95±0.08 | 0.90±0.08 | 0.80±0.07 | 0.86±0.09 | 0.80±0.09 |
| G&S.cingul-Mid-Ant | 0.72±0.11 | 0.75±0.08 | 0.67±0.10 | 0.62±0.07 | 0.63±0.07 | 0.54±0.06 | 0.57±0.06 | 0.61±0.08 | 0.58±0.06 |
| G&S.cingul-Mid-Post | 0.75±0.11 | 0.83±0.09 | 0.77±0.10 | 0.65±0.08 | 0.69±0.08 | 0.62±0.07 | 0.59±0.07 | 0.65±0.08 | 0.65±0.07 |
| G.cingul-Post-dorsal | 0.77±0.10 | **0.88±0.08** | **0.90±0.10** | 0.82±0.09 | 0.89±0.09 | 0.84±0.09 | 0.72±0.09 | **0.85±0.09** | 0.82±0.10 |
| G.cingul-Post-ventral | 0.66±0.10 | 0.72±0.08 | 0.68±0.10 | 0.57±0.09 | 0.64±0.08 | 0.53±0.07 | 0.50±0.08 | **0.62±0.08** | 0.54±0.07 |
| G.cuneus | 0.66±0.09 | 0.71±0.07 | 0.63±0.09 | 0.61±0.09 | 0.68±0.07 | 0.52±0.07 | 0.52±0.08 | **0.65±0.07** | 0.55±0.07 |
| G.front_inf-Opercular | 0.68±0.10 | 0.73±0.08 | 0.69±0.10 | 0.53±0.07 | 0.54±0.07 | 0.47±0.08 | 0.54±0.06 | 0.54±0.08 | 0.53±0.07 |
| G.front_inf-Orbital | 0.75±0.08 | **0.88±0.07** | 0.81±0.08 | 0.80±0.07 | 0.79±0.07 | 0.66±0.08 | 0.74±0.07 | 0.75±0.07 | 0.71±0.07 |
| G.front_inf-Triangul | 0.81±0.10 | 0.89±0.07 | 0.84±0.10 | 0.75±0.08 | 0.74±0.07 | 0.62±0.08 | 0.74±0.07 | 0.72±0.07 | 0.67±0.08 |
| G.front_middle | 1.11±0.13 | 1.17±0.09 | 1.15±0.13 | 1.00±0.10 | 1.08±0.09 | 0.99±0.11 | 1.01±0.10 | 1.05±0.09 | 1.01±0.10 |
| G.front_sup | 1.22±0.12 | 1.30±0.10 | 1.28±0.11 | 1.17±0.09 | 1.23±0.08 | 1.17±0.09 | 1.07±0.09 | 1.15±0.09 | 1.14±0.09 |
| G.Ins.lg&S.cent.ins | 0.57±0.09 | 0.59±0.07 | 0.55±0.08 | 0.48±0.07 | 0.49±0.06 | 0.42±0.07 | 0.46±0.06 | 0.51±0.07 | 0.48±0.07 |
| G.insular_short | 0.54±0.08 | 0.60±0.06 | 0.53±0.08 | 0.49±0.05 | 0.49±0.06 | 0.43±0.07 | 0.51±0.05 | 0.50±0.07 | 0.50±0.06 |
| G.occipital_middle | 0.83±0.10 | 0.90±0.09 | 0.86±0.10 | 0.68±0.09 | 0.73±0.08 | 0.62±0.07 | 0.60±0.08 | 0.68±0.08 | 0.61±0.06 |
| G.occipital_sup | 0.71±0.09 | 0.79±0.08 | 0.71±0.09 | 0.60±0.09 | 0.66±0.07 | 0.53±0.07 | 0.52±0.08 | 0.64±0.07 | 0.54±0.07 |
| G.oc-temp_lat-fusifor | 0.82±0.10 | 0.86±0.09 | 0.84±0.11 | 0.68±0.09 | 0.75±0.08 | 0.63±0.08 | 0.62±0.07 | 0.69±0.08 | 0.63±0.07 |
| G.oc-temp_med-Lingual | 0.69±0.08 | 0.72±0.08 | 0.67±0.10 | 0.62±0.09 | 0.70±0.07 | 0.56±0.08 | 0.55±0.08 | **0.68±0.07** | 0.58±0.07 |
| G.oc-temp_med-Parahip | 0.52±0.08 | 0.57±0.08 | 0.51±0.08 | 0.49±0.07 | 0.55±0.08 | 0.48±0.08 | 0.42±0.06 | 0.46±0.07 | 0.41±0.08 |
| G.orbital | 1.04±0.09 | 1.07±0.07 | 1.06±0.09 | 1.01±0.08 | 1.02±0.08 | 0.89±0.09 | 0.93±0.08 | 0.94±0.08 | 0.87±0.08 |
| G.pariet_inf-Angular | 0.87±0.10 | **0.99±0.08** | 0.94±0.10 | 0.79±0.08 | 0.88±0.08 | 0.79±0.09 | 0.74±0.07 | 0.82±0.08 | 0.78±0.08 |
| G.pariet_inf-Supramar | 0.75±0.09 | 0.82±0.08 | 0.78±0.10 | 0.55±0.07 | 0.62±0.07 | 0.55±0.08 | 0.54±0.07 | 0.60±0.08 | 0.59±0.08 |
| G.parietal_sup | 0.84±0.11 | 0.94±0.09 | 0.89±0.10 | 0.67±0.11 | **0.78±0.08** | 0.69±0.07 | 0.59±0.09 | **0.73±0.09** | 0.67±0.08 |
| G.postcentral | 0.66±0.09 | **0.76±0.07** | 0.71±0.09 | 0.39±0.08 | **0.51±0.07** | 0.47±0.07 | 0.37±0.07 | **0.50±0.08** | 0.50±0.07 |
| G.precentral | 0.72±0.09 | 0.81±0.07 | 0.75±0.09 | 0.50±0.08 | **0.62±0.07** | 0.54±0.07 | 0.48±0.07 | **0.59±0.08** | 0.56±0.08 |
| G.precuneus | 0.86±0.12 | **0.97±0.09** | 0.93±0.11 | 0.78±0.11 | 0.87±0.09 | 0.77±0.08 | 0.69±0.09 | **0.82±0.09** | 0.75±0.09 |
| G.rectus | 0.53±0.07 | 0.54±0.07 | 0.56±0.09 | 0.37±0.06 | 0.41±0.08 | 0.46±0.10 | 0.23±0.07 | 0.27±0.07 | 0.30±0.08 |
| G.subcallosal | 0.45±0.06 | 0.44±0.06 | 0.38±0.08 | 0.42±0.05 | 0.41±0.06 | 0.37±0.07 | 0.21±0.05 | 0.23±0.07 | 0.17±0.08 |
| G.temp_sup-G.T.transv | 0.54±0.10 | 0.58±0.07 | 0.57±0.09 | 0.41±0.06 | 0.41±0.07 | 0.38±0.07 | 0.42±0.06 | 0.46±0.08 | 0.44±0.08 |
| G.temp_sup-Lateral | 0.80±0.10 | 0.91±0.09 | 0.86±0.11 | 0.76±0.08 | 0.79±0.08 | 0.70±0.08 | 0.69±0.07 | 0.73±0.09 | 0.71±0.09 |
| G.temp_sup-Plan.polar | 0.48±0.08 | **0.59±0.07** | 0.54±0.08 | 0.52±0.07 | 0.55±0.07 | 0.47±0.08 | 0.50±0.07 | 0.50±0.08 | 0.47±0.09 |
| G.temp_sup-Plan-tempo | 0.71±0.09 | 0.77±0.08 | 0.72±0.10 | 0.53±0.07 | 0.56±0.07 | 0.48±0.07 | 0.49±0.06 | 0.55±0.08 | 0.52±0.07 |
| G.temporal_inf | 0.84±0.12 | 0.90±0.09 | 0.87±0.11 | 0.73±0.09 | 0.81±0.08 | 0.72±0.09 | 0.64±0.08 | 0.70±0.08 | 0.65±0.09 |
| G.temporal_middle | 0.90±0.11 | 0.99±0.09 | 0.95±0.10 | 0.86±0.09 | 0.89±0.08 | 0.79±0.10 | 0.74±0.07 | 0.79±0.08 | 0.73±0.10 |
| Lat.Fis-ant-Horizont | 0.58±0.08 | **0.69±0.06** | 0.62±0.08 | 0.63±0.06 | 0.60±0.07 | 0.49±0.08 | 0.63±0.07 | 0.61±0.07 | 0.59±0.08 |
| Lat.Fis-ant-Vertical | 0.55±0.08 | 0.64±0.06 | 0.59±0.08 | 0.57±0.06 | 0.53±0.06 | 0.44±0.07 | 0.58±0.07 | 0.53±0.07 | 0.52±0.07 |
| Lat.Fis-post | 0.62±0.09 | 0.64±0.07 | 0.61±0.10 | 0.43±0.07 | 0.44±0.07 | 0.40±0.08 | 0.41±0.06 | 0.47±0.08 | 0.46±0.07 |
| Medial.wall | 0.69±0.10 | 0.78±0.08 | 0.73±0.09 | 0.74±0.08 | 0.75±0.09 | 0.69±0.09 | 0.54±0.08 | 0.62±0.08 | 0.56±0.09 |
| Pole.occipital | 0.68±0.08 | 0.69±0.07 | 0.66±0.09 | 0.65±0.08 | 0.72±0.07 | 0.58±0.09 | 0.57±0.08 | **0.70±0.08** | 0.59±0.07 |
| Pole.temporal | 0.56±0.09 | 0.61±0.07 | 0.49±0.08 | 0.57±0.07 | 0.61±0.08 | 0.51±0.09 | 0.48±0.07 | 0.46±0.07 | 0.41±0.09 |
| S.calcarine | 0.68±0.09 | 0.73±0.08 | 0.62±0.09 | 0.54±0.08 | 0.60±0.07 | 0.46±0.07 | 0.46±0.07 | **0.58±0.07** | 0.47±0.07 |
| S.central | 0.66±0.09 | 0.75±0.07 | 0.69±0.09 | 0.41±0.08 | 0.52±0.07 | 0.47±0.07 | 0.39±0.07 | **0.51±0.08** | 0.51±0.07 |
| S.cingul-Marginalis | 0.78±0.11 | 0.84±0.09 | 0.81±0.10 | 0.68±0.09 | 0.75±0.09 | 0.70±0.07 | 0.64±0.08 | 0.73±0.09 | 0.70±0.08 |
| S.circular.insula_ant | 0.58±0.08 | 0.66±0.07 | 0.59±0.08 | 0.67±0.05 | 0.62±0.06 | 0.59±0.06 | 0.65±0.06 | 0.65±0.07 | 0.64±0.08 |
| S.circular.insula_inf | 0.47±0.08 | 0.53±0.07 | 0.50±0.08 | 0.47±0.07 | 0.46±0.06 | 0.42±0.07 | 0.46±0.06 | 0.52±0.08 | 0.50±0.08 |
| S.circular.insula_sup | 0.47±0.08 | 0.52±0.06 | 0.45±0.08 | 0.30±0.05 | 0.32±0.06 | 0.26±0.06 | 0.35±0.05 | 0.36±0.06 | 0.36±0.05 |
| S.collat.transv_ant | 0.56±0.11 | 0.66±0.08 | 0.60±0.09 | 0.49±0.08 | 0.57±0.07 | 0.51±0.07 | 0.43±0.06 | 0.44±0.06 | 0.41±0.07 |
| S.collat.transv_post | 0.76±0.08 | 0.78±0.09 | 0.74±0.10 | 0.66±0.09 | 0.71±0.08 | 0.57±0.08 | 0.58±0.08 | 0.67±0.08 | 0.57±0.07 |
| S.front_inf | 0.81±0.10 | 0.89±0.08 | 0.85±0.11 | 0.69±0.09 | 0.75±0.08 | 0.63±0.09 | 0.69±0.08 | 0.72±0.08 | 0.67±0.08 |
| S.front_middle | 1.18±0.13 | 1.24±0.10 | 1.22±0.14 | 1.13±0.11 | 1.18±0.11 | 1.14±0.13 | 1.16±0.12 | 1.20±0.11 | 1.20±0.12 |
| S.front_sup | 0.93±0.12 | 1.01±0.10 | 0.98±0.12 | 0.88±0.09 | 0.99±0.09 | 0.93±0.10 | 0.87±0.09 | **0.96±0.10** | 0.92±0.10 |
| S.interm.prim-Jensen | 0.76±0.10 | 0.83±0.08 | 0.79±0.11 | 0.66±0.08 | 0.70±0.08 | 0.60±0.08 | 0.64±0.08 | 0.70±0.09 | 0.60±0.08 |
| S.intrapariet&P.trans | 0.79±0.10 | 0.88±0.08 | 0.85±0.10 | 0.61±0.09 | 0.71±0.08 | 0.64±0.07 | 0.57±0.09 | 0.68±0.08 | 0.63±0.07 |
| S.oc.middle&Lunatus | 0.76±0.10 | 0.83±0.09 | 0.77±0.11 | 0.60±0.08 | 0.64±0.08 | 0.52±0.07 | 0.52±0.07 | 0.60±0.08 | 0.51±0.06 |
| S.oc.sup&transversal | 0.69±0.09 | 0.78±0.09 | 0.71±0.10 | 0.54±0.09 | 0.57±0.07 | 0.45±0.06 | 0.47±0.07 | 0.55±0.07 | 0.45±0.05 |
| S.occipital_ant | 0.71±0.10 | 0.79±0.09 | 0.76±0.10 | 0.54±0.08 | 0.58±0.07 | 0.49±0.06 | 0.48±0.08 | 0.55±0.08 | 0.51±0.05 |
| S.oc-temp_lat | 0.76±0.11 | 0.86±0.09 | 0.84±0.10 | 0.59±0.09 | 0.68±0.07 | 0.59±0.07 | 0.55±0.07 | 0.60±0.07 | 0.59±0.07 |
| S.oc-temp_med&Lingual | 0.70±0.09 | 0.79±0.09 | 0.73±0.09 | 0.56±0.08 | 0.60±0.07 | 0.50±0.06 | 0.49±0.07 | 0.58±0.08 | 0.48±0.06 |
| S.orbital.lateral | 0.83±0.08 | 0.91±0.07 | 0.86±0.09 | 0.78±0.06 | 0.81±0.06 | 0.71±0.09 | 0.75±0.08 | 0.80±0.07 | 0.76±0.09 |
| S.orbital.med-olfact | 0.50±0.07 | 0.49±0.06 | 0.51±0.08 | 0.37±0.06 | 0.40±0.07 | 0.37±0.08 | 0.22±0.05 | 0.28±0.07 | 0.23±0.08 |
| S.orbital-H.Shaped | 0.83±0.11 | 0.84±0.08 | 0.81±0.09 | 0.77±0.09 | 0.78±0.09 | 0.67±0.08 | 0.70±0.08 | 0.72±0.07 | 0.63±0.08 |
| S.parieto.occipital | 0.71±0.10 | 0.79±0.08 | 0.69±0.09 | 0.58±0.09 | 0.63±0.08 | 0.51±0.06 | 0.51±0.08 | **0.63±0.07** | 0.52±0.06 |
| S.pericallosal | 0.77±0.10 | 0.82±0.08 | 0.81±0.09 | 0.79±0.08 | 0.81±0.08 | 0.73±0.07 | 0.67±0.08 | **0.78±0.08** | 0.72±0.07 |
| S.postcentral | 0.67±0.09 | 0.76±0.07 | 0.71±0.09 | 0.38±0.08 | **0.50±0.08** | 0.46±0.07 | 0.37±0.08 | **0.49±0.08** | 0.49±0.07 |
| S.precentral-inf-part | 0.71±0.10 | 0.80±0.08 | 0.74±0.09 | 0.56±0.09 | 0.64±0.06 | 0.53±0.07 | 0.55±0.07 | 0.60±0.07 | 0.55±0.07 |
| S.precentral-sup-part | 0.70±0.10 | 0.80±0.08 | 0.76±0.09 | 0.54±0.09 | 0.64±0.07 | 0.56±0.06 | 0.49±0.08 | **0.61±0.08** | 0.58±0.07 |
| S.suborbital | 0.71±0.07 | 0.75±0.07 | 0.77±0.09 | 0.61±0.08 | 0.62±0.09 | 0.61±0.10 | 0.41±0.08 | 0.47±0.08 | 0.49±0.09 |
| S.subparietal | 0.80±0.11 | **0.92±0.09** | 0.89±0.11 | 0.78±0.09 | 0.87±0.09 | 0.80±0.08 | 0.72±0.10 | **0.85±0.09** | 0.79±0.09 |
| S.temporal_inf | 0.80±0.10 | 0.90±0.09 | 0.83±0.10 | 0.69±0.09 | 0.77±0.09 | 0.68±0.09 | 0.58±0.07 | 0.64±0.08 | 0.57±0.09 |
| S.temporal_sup | 0.87±0.09 | **0.98±0.08** | 0.94±0.10 | 0.87±0.08 | 0.89±0.08 | 0.80±0.08 | 0.79±0.08 | 0.84±0.08 | 0.80±0.09 |
| S.temporal_transverse | 0.58±0.10 | 0.64±0.08 | 0.62±0.10 | 0.45±0.07 | 0.46±0.07 | 0.42±0.07 | 0.44±0.06 | 0.49±0.08 | 0.48±0.08 |

